# A YIPF5-GOT1A/B complex directs a transcription-independent function of ATF6 in ER export

**DOI:** 10.1101/2023.12.12.569033

**Authors:** Paul Cramer, Yoji Yonemura, Laura Behrendt, Aleksandra Marszalek, Mara Sannai, William Durso, Cagatay Günes, Karol Szafranski, Nobuhiro Nakamura, Tornike Nasrashvili, Johanna Mayer, Björn von Eyss, Christoph Kaether

**Affiliations:** Leibniz Institute on Aging-Fritz Lipmann Institute, 07745 Jena, Germany; Faculty of Life Sciences, Kyoto Sangyo University, Motoyama, Kamigamo, Kita, Kyoto 603-8555, Japan; Department of Neurobiology, Care Sciences and Society, Karolinska Institute, 17177 Stockholm, Sweden; Laboratory of Ion Channel Pathophysiology, Faculty of Brain Pathology, Doshisha University, Kyoto, Japan; Department of Urology, Ulm University Hospital, Ulm, Germany

**Author notes:** CK and BvE should be considered joint senior authors. **Correspondence should be addressed to:** Christoph Kaether, Leibniz Institute on Aging, Fritz-Lipmann Institute, Beutenbergstr. 11, 07745 Jena, Germany, Björn von Eyss, Leibniz Institute on Aging, Fritz-Lipmann Institute, Beutenbergstr. 11, 07745 Jena, Germany.

**Keywords:** endoplasmic reticulum, ER exit sites, COPII, secretory transport, ER stress, ATF6

## Abstract

Exit from the endoplasmic reticulum is mediated by the Sar1/COPII machinery and a number of accessory factors. How the initial steps of cargo recruitment upstream of Sar1/COPII are mediated remains unclear, but the dihydropyridine FLI-06 inhibits cargo recruitment into ER exit sites. Here, we used chemical genetics screening approaches in conjunction with FLI-06 treatment and identified the ER membrane proteins YIPF5 and GOT1A/B as putative components of early export processes. Surprisingly, the two homologous proteins GOT1A and GOT1B, coded by *GOLT1A* and *GOLT1B*, respectively, exhibited opposite functions after treatment with FLI-06: increasing the expression of GOT1A or reducing the expression of GOT1B or YIPF5 prevented inhibition of ER-export by FLI-06. Inhibiting ER export with FLI-06 elicited a specific ER stress-related gene expression signature distinct from the ER-stress signature induced by Thapsigargin. The interactomes of GOT1A and GOT1B suggested a connection to ER-stress mediators. Moreover, RNA-Seq data showed that FLI-06-induced genes are strongly enriched for ATF6 target genes which are suppressed by GOLT1A overexpression or GOLT1B knock-down. This suggests that ATF6 signaling is involved in FLI-06-mediated toxicity, and we could demonstrate that siRNA-mediated knock-down or specific inhibitor of ATF6 rescued cells from FLI-06-mediated cell death. Knock-down or inhibition of ATF6 is sufficient to resume transport from the ER under FLI-06-treatment, suggesting that ATF6 is directly involved in the FLI-06-mediated ER-export block. Surprisingly, our data show that this ATF6 function is independent of *de novo* transcription, implying a novel, transcription-independent function of ATF6.

## Introduction

The secretory pathway is fundamental in all eukaryotic cells, with about one third of all proteins being directly or indirectly connected to it. The largest organelle of the secretory pathway is the endoplasmic reticulum (ER), the site of synthesis for most secreted and for most membrane proteins. Export from the ER occurs at ER exit sites (ERES), specialized domains at the ER where export cargo is concentrated (reviewed in (Aridor, 2018, Barlowe & Helenius, 2016, Zanetti et al., 2012)). From here the cargo is transported to the Golgi, involving the ER-Golgi intermediate compartment (ERGIC) (Schweizer et al., 1994). Export out of the ER is mediated by Sar1, a small GTPase, and the COPII coat complex that consists of the inner coat proteins Sec23/24 and the outer coat proteins Sec13/31 (reviewed in (Zanetti et al., 2012)). This is the basic machinery shown *in vitro* to bud off vesicles containing cargo from ER membranes. In mammalian cells the COPII machinery likely coats the neck of nascent ERES and is not leaving them (Malis et al., 2022, Shomron et al., 2021, Weigel et al., 2021). Additional proteins like Sec12, Sec16, cornichons, ERGIC53, p24 family member and more are needed for efficient concentration and export of cargo. Some of these proteins are needed for all kind of cargoes, others only for specialized cargo like huge collagen fibers or scales (reviewed in (Barlowe & Helenius, 2016, Gomez-Navarro & Miller, 2016, McCaughey & Stephens, 2018, Raote et al., 2023)).

Some proteins were identified in the early secretory pathway with ill-defined or controversial roles in ER export, among them YIPF5 and GOT1B. YIPF5 (aliases FinGER5, YIP1A, YIP1, Yip1p in yeast) is a small five-span transmembrane domain (TMD) protein of the ER and the early secretory pathway. A role for YIPF5 in COPII vesicle biogenesis in yeast was suggested (Heidtman et al., 2003) and in HeLa cells it interacts with Sec23/24 (Tang et al., 2001), but knockdown in mammalian cells did not affect secretion or viability (Kranjc et al., 2017, Yoshida et al., 2008). YIPF5 is in a complex with YIF1A (Yoshida et al., 2008) and interacts with GOT1B (Hein et al., 2015). GOT1B (aliases: GOT1, Got1p in yeast), the protein encoded by the *GOLT1B* gene, is a four-span TMD protein of the early secretory pathway. Like YIPF5, GOT1B is a component of COPI and COPII vesicles (Adolf et al., 2019) and, in rice, interacts with the COPII protein Sec23 (Wang et al., 2016). Interestingly, in yeast overexpression of Got1p rescues a YIPF5-thermosensitive allele, demonstrating a genetic interaction (Lorente-Rodriguez et al., 2009). GOT1A, encoded by *GOLT1A*, is a highly conserved homologue of GOT1B (Conchon et al., 1999), with 54% identity and 89% similarity on the protein level. According to protein atlas (https://www.proteinatlas.org/) expression is high in liver, pituitary and gastrointestinal tract. Little is known about its molecular function, but proliferation of various tumors seems to depend on GOT1A expression (Ikeda et al., 2015, Zhang et al., 2017) Recently, evidence was presented that COPII has additional functions in controlling the access to ERES (Ma et al., 2017). In addition, there seem to be as yet unidentified processes upstream of Sar1/COPII that recruit cargo to ERES. These processes are inhibited by the secretion inhibitor FLI-06, a dihydropyridine inhibiting ER export of all cargos tested to date (Krämer et al., 2013, Yonemura et al., 2016).

Upon disturbances in ER proteostasis like accumulation of unfolded or folded proteins (ER stress), the ER activates a stereotypic response to mitigate ER stress, the unfolded protein response (UPR). Three arms of the UPR, mediated by inositol-requiring protein 1 (IRE1), activating transcription factor-6 (ATF6) and protein kinase RNA (PKR)-like ER kinase (PERK) re-balance proteostasis in the ER. This is achieved by for example up-regulating the expression of ER-chaperones, inducing ER-associated degradation (ERAD) and reducing ER-import (for review seeHetz et al., 2020, Ron & Walter, 2007). If proteostasis cannot be restored, apoptosis is initiated (for review seeHetz et al., 2020, Ron & Walter, 2007). The UPR also increases ER export capacity by up-regulating expression of genes involved in ER-Golgi transport (Higashio & Kohno, 2002, Sato et al., 2002, Shaffer et al., 2004, Sriburi et al., 2007). Complex signaling pathways link ER import and folding capacity on one hand with ER export capacity on the other hand (Centonze et al., 2019, Subramanian et al., 2019). It is still an open question whether there are mechanisms in place to down-regulate or shut down ER export completely when import into the ER and folding of nascent proteins are minimized.

Using pharmacogenomic screening approaches, we here show that YIPF5, GOT1A and GOT1B are involved in a FLI-06-susceptible ER export step. FLI-06 stabilizes ATF6 full-length protein in the ER, which contributes but is not sufficient for the secretion block. This suggests a novel role for ATF6 beyond its well-established role as a transcription factor in UPR.

## Results

### A chemical-genetic approach to identify pathways inhibited by secretion inhibitors

CRISPR/Cas-based loss-of function (LOF) and gain-of-function (GOF) screens are powerful tools to identify putative targets and mechanisms of small compounds and to identify novel regulatory pathways (Konermann et al., 2015, Shalem et al., 2014) (Elster et al., 2018). Here, we wanted to test whether a similar approach could be used in conjunction with small compounds targeting specific steps in the secretory pathway. In this way, we aimed to identify genetic perturbations that allow cells to survive treatment with inhibitors targeting the secretory pathway (Fig. 1a,b). We first tested whether this approach could be addressed using the SAM (synergistic activation mediator) system. SAM is an adaptation of CRISPR_activation_, in combination with a genome-wide lentiviral sgRNA library (Konermann et al., 2015). The SAM system consists of dCas9-VP64 and MCP-p65-HSF1 fusion proteins, which are recruited to the promoter of the sgRNA-targeted gene, leading to further recruitment of the transcription machinery and overexpression of the endogenous locus (Konermann et al., 2015). We performed proof-of-concept experiments for such a GOF screen with a well-characterized inhibitor of secretion, Brefeldin A (BFA). BFA is a widely used small compound that efficiently blocks early secretion and leads to a fusion of the Golgi with the ER ((Fujiwara et al., 1988, Klausner et al., 1992, Misumi et al., 1986). BFA has three main targets, GBF1 and BIG1/2, all of which are guanine nucleotide exchange factors (ArfGEFS) involved in the recruitment of coat proteins (Claude et al., 1999, Yamaji et al., 2000). GBF1 is the target of BFA responsible for the secretion inhibition at the early secretory pathway (Saenz et al., 2009).

**Figure 1.**
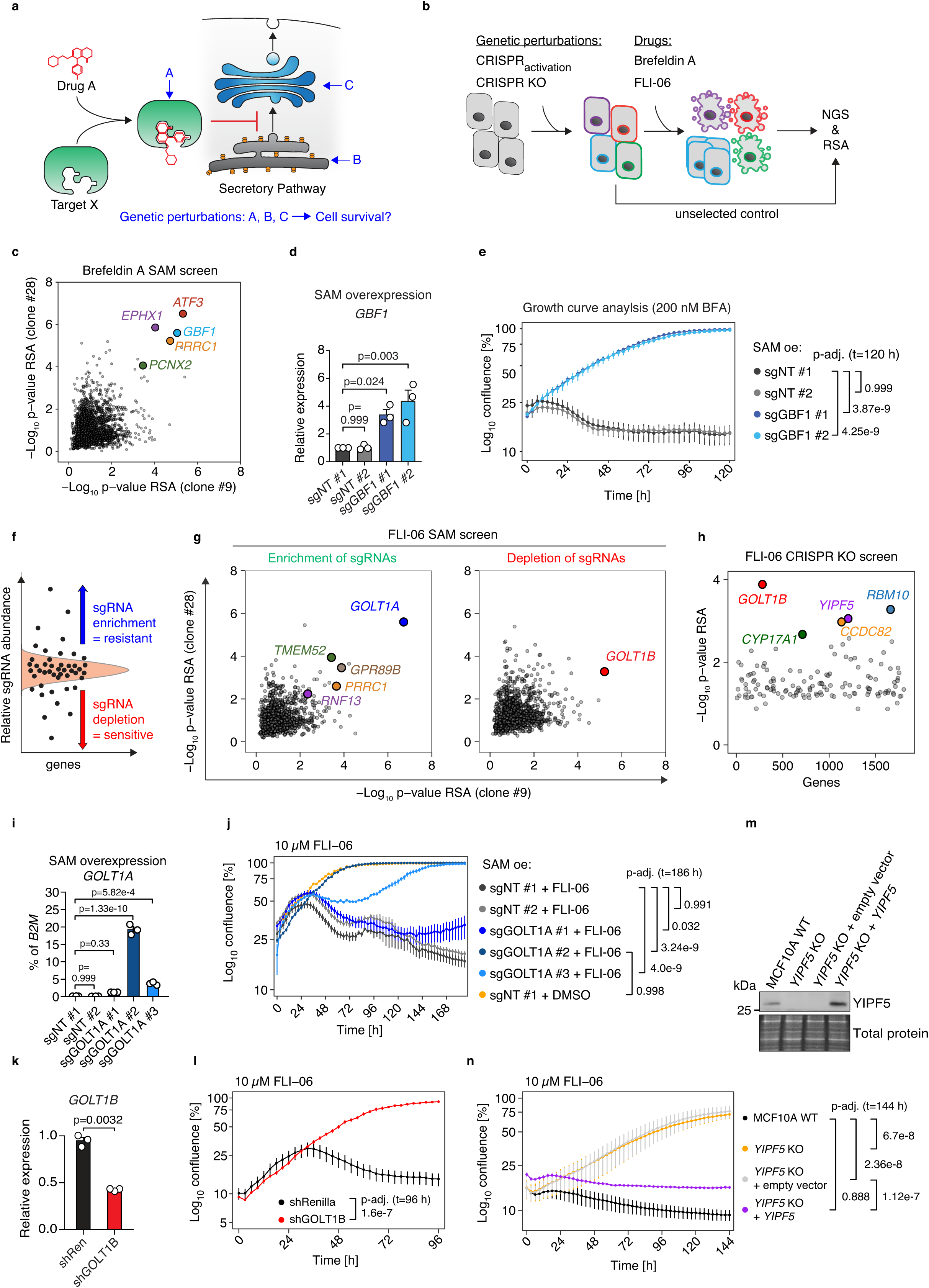
Genome-wide CRISPR screens identify *GOLT1A/B* and *YIPF5* as modulators of FLI-06-dependent phenotypes. **a)** Schematic of pharmacogenetic screens to identify factors that modulate the effect of secretion inhibitors. **b)** General procedure of CRISPR_activation_ and CRISPR knock-out (KO) screening approaches. MCF10A cells were infected with the genome-wide lentiviral SAM (synergistic activation mediator) or GeCKO v2 sgRNA library. After selection with Brefeldin A (BFA) or FLI-06, the sgRNA distribution was determined by NGS (next generation sequencing) and analyzed using the RSA (redundant siRNA activity) algorithm in comparison with unselected control cells. **c)** CRISPR_activation_ screen for Brefeldin A (BFA, 100 nM) resistance in clonal MCF10A cells using the SAM system. Displayed is the RSA analysis for sgRNA enrichment from two MCF10A-SAM clones (#9 and #28). Each dot represents one gene, gene names of selected hits are shown. **d-e**) Validation of GBF1 as BFA target. qPCR expression analysis (**d**) of *GBF1* and IncuCyte growth curve analysis upon BFA treatment (**e**) in MCF10A-SAM cells stably expressing sgGBF1 or non-targeting (NT) sgRNAs. Displayed is the confluency over time. **f**) Principle to identify genetic perturbations that confer resistance or sensitivity to secretion inhibitors by sgRNA enrichment or depletion, respectively. **g**) CRISPR_a_ screen for FLI-06 (10 µM) resistance in clonal MCF10A cells (#9 and #28) using the SAM system. RSA analysis for both clones was performed for sgRNA enrichment (left) or sgRNA depletion (right). Each dot represents one gene. **h**) CRISPR-KO screen for FLI-06 (10 µM) resistance in human BJ-Tert cells using the GeCKO sgRNA library. RSA analysis was performed for sgRNA enrichment. Each dot represents one gene. **i-n**) Validation of *GOLT1A*, *GOLT1B* and *YIPF5* as mediators of resistance/sensitivity to FLI-06. qPCR expression analysis (**i**) of *GOLT1A* and IncuCyte growth curve analysis upon FLI-06 (10 µM) treatment (**j**) in MCF10A-SAM clone #9 stably expressing sgGOLT1A or non-targeting sgRNAs. qPCR expression analysis (**k**) of *GOLT1B* and IncuCyte growth curve analysis upon FLI-06 treatment (**l**) in MCF10A cells stably expressing shGOLT1B or shRenilla as control. **m**) Immunoblotting analysis with total protein as loading control and **n**) IncuCyte growth curve analysis upon FLI-06 treatment in *YIPF5* KO and *YIPF5* or empty vector re-expressing MCF10A cells. For qPCR, *B2M* was used as housekeeping gene. Data are shown as mean of three biological replicates and error bars represent SEM (**d, e, i, j, k, l, n**). Statistical testing: one-way-ANOVA test with Tukey HSD post-hoc test (**d, e, i, j, l, n**) and unpaired two-sided Welch t-test (**k**).

For GOF SAM-screening (Fig. 1), we used two independent MCF10A cell clones stably overexpressing the required components of the SAM system, dCas9-VP64 and MCP-p65-HSF1. These cells were subsequently infected with a genome-wide lentiviral SAM sgRNA library and cultured for 21 days in the presence of 100 nM BFA, a concentration that normally results in cell death of all cells (Fig. S1a-g). After next generation sequencing (NGS) of the sgRNA cassette, we compared the sgRNA frequencies before and after treatment (schematized in Fig 1b). Subsequently, we integrated the values from different sgRNAs targeting the same gene by redundant siRNA activity (RSA)(Konig et al., 2007). As expected, one of the top hits in both MCF10A-SAM clones was *GBF1* (Fig. 1c, S1a,b), demonstrating the power of GOF screens to identify targets of secretion inhibitors.

Confirming the screening results, SAM-mediated overexpression of *GBF1* by individual sgRNAs led to a successful upregulation of the endogenous gene (Fig. 1d) resulting in resistance to BFA up to a concentration of 200 nM BFA (Fig. 1e). Apart from *GBF1*, we could identify *ATF3*, *EPHX1*, *PRRC1* and *PCNX2* as previously unknown mediators of BFA resistance (Fig. S1c-h).

Taken together, these results demonstrate the power of SAM screening to identify genes involved in early secretion.

### Deletion of GOLT1B and YIPF5 or overexpression of GOLT1A mediate resistance to FLI-06

To identify unknown components of the selection/concentration mechanisms upstream of COPII, we next combined SAM screening with the secretion inhibitor FLI-06 (Fig. 1f-n, Fig. S1i-o), a molecule that blocks ER-exit (Krämer et al., 2013, Yonemura et al., 2016). The FLI-06-mediated secretion block leads to growth arrest and eventually cell death after two to three days (Fig. 1j, sgNT#1,2). Similar to the BFA screen, we next performed a SAM screen with FLI-06 (Fig. 1f,g). After infection with the sgRNA library, cells were cultured in the presence of 10 µM FLI-06 for 21 days. We determined 10 µM FLI-06 as a concentration with moderate stringency to identify genes that mediate resistance or hypersensitivity upon their overexpression, respectively (Fig. 1f). After screening and NGS, RSA analyses identified multiple genes that either conferred resistance or hypersensitivity to FLI-06 (Fig. 1f,g). By far the most significant hit conferring FLI-06 resistance was *GOLT1A*, encoding the protein GOT1A (Fig. 1g). Surprisingly, *GOLT1B*, encoding GOT1B, was among the top hits that led to FLI-06 hypersensitivity when overexpressed (Fig. 1g). This was unexpected since GOT1A and GOT1B are highly conserved paralogues (Conchon et al., 1999) (Fig. S2a). Little is known about the molecular function of GOT1A and GOT1B, but our results imply opposing functions for these paralogues in relation to FLI-06.

To validate GOT1A as hit of the GOF screen, we infected MCF10A SAM cells with individual sgRNAs targeting *GOLT1A*. The different sgRNAs induced *GOLT1A* to a varying degree (Fig. 1i). *GOLT1A* expression consistently correlated with FLI-06 resistance and the most potent sgRNA completely restored cell proliferation under FLI-06 treatment to the level of DMSO-treated cells (Fig. 1j). Notably, *GOLT1A* overexpression still rescued FLI-06 sensitivity at concentrations as high as 20 µM (Fig. S2b). Additionally, sgRNA-mediated *GOLT1A* overexpression did not affect cell proliferation in the absence of FLI-06 (Supp. Fig. S2c). Endogenous *GOLT1A* mRNA was barely detectable in MCF10A cells, resulting in a sgRNA-mediated absolute expression level of *GOLT1A* mRNA in the range of endogenous *GOLT1B* expression (compare Fig. 1i and S2d). Similarly, overexpression of the other hits *TMEM52* and *RNF13* also conferred FLI-06 resistance, but to a smaller extent than *GOLT1A*, while *PRRC1* and *GPR89B* could not be validated (Fig S1k-o).

In the screen, SAM-activation of *GOLT1B* rendered cells hypersensitive to FLI-06. To validate this finding, cells expressing individual sgRNAs were treated with 5 μM FLI-06 and their viability assayed (Fig. S2d,e). sgRNAs #1 and #3, which both increase expression of GOLT1B, but not ineffective sgRNA #2 mediated hypersensitivity to FLI-06 (Fig. S2d,e).

To complement the GOF screen and to validate our previous findings in another cell line, we next performed a LOF screen in primary human BJ cells (Fig. 1h). To this end, we infected BJ cells with a lentiviral genome-wide CRISPR KO library (Shalem et al., 2014) and treated them with 10 µM FLI-06. After several weeks of selection, genomic DNA was extracted and subsequent analyses were performed similarly to the SAM screen. Strikingly, *GOLT1B* was the top hit together with *YIPF5* (Fig. 1h), the gene encoding for YIPF5 - a known GOT1B-interacting protein (Hein et al., 2015). We validated GOT1B and YIPF5 as mediators of FLI-06 sensitivity. Stable knock-down of *GOLT1B* in MCF10A cells by shRNAs rendered cells resistant to FLI-06 (Fig. 1 k, l), as did CRISPR/Cas-mediated knock-out of *YIPF5* (Fig. 1 m,n). Stable re-expression of YIPF5, but not empty vector, re-sensitized cells to FLI-06 (Fig. 1 m,n).

Taken together, these complementary genome-wide screening results identify GOT1B and its interaction partner YIPF5 as factors that mediate sensitivity to FLI-06 and GOT1A as a factor that mediates resistance to FLI-06 when overexpressed.

### YIPF5, GOT1A and GOT1B are proteins involved in an FLI-06 sensitive step of ER export

Given the significant enrichment of GOT1A and GOT1B in our screens and their unexpected opposing roles during FLI-06 treatment, we focused on these two proteins to analyze the mechanistic basis for their role in FLI-06 resistance. Furthermore, GOT1B and its yeast homologue Got1p have been linked to the secretory pathway (Adolf et al., 2019, Conchon et al., 1999), but its homologue GOT1A has not been studied on the molecular level so far.

FLI-06 treatment results in Golgi dispersal after 3-4 hours of treatment (Krämer et al., 2013). MCF10A cells overexpressing endogenous *GOLT1A* by SAM (Fig. S2b,c,f,g) or Flag-*GOLT1A* cDNA (Fig. 2a,b, Fig S2h,j) were completely resistant to FLI-06-mediated Golgi dispersal and cell death. In contrast, MCF10A cells overexpressing *GOLT1B*-Flag (Fig. 2a,b) or a sgRNA activating endogenous *GOLT1B* expression (Fig. S2d-g) demonstrated an increased Golgi dispersal. Similarly, *YIPF5* knockout cells did not show Golgi dispersal after FLI-06 treatment but reconstitution with YIPF5 restored this effect (Fig. 2c,d). In order to analyze whether GOT1A/B and YIPF5 impact on the FLI-06-mediated secretion block, we performed VSVG-EGFP temperature-shift assays in HeLa cells (Fig. 3a). In this assay, transport of a temperature-sensitive mutant (tsO45) of the VSVG-protein (Gallione & Rose, 1985) can be assayed by fluorescence microscopy (Fig. 3b) or by assaying N-glycosylation status with endoglycosidase H (endoH’, Fig. 3d). FLI-06 blocks this transport at the ER (Krämer et al., 2013, Yonemura et al., 2016), Fig. 3b). Overexpression of HA-tagged GOT1A or depletion of GOT1B could largely restore ER export after FLI-06 treatment (Fig. 3b-e). Knockdown of YIPF5, however, only had a weak effect in this assay which did not reach significance, most likely due to the poor YIPF5 knockdown efficiency (Fig. 3c, e).

**Fig. 2:**
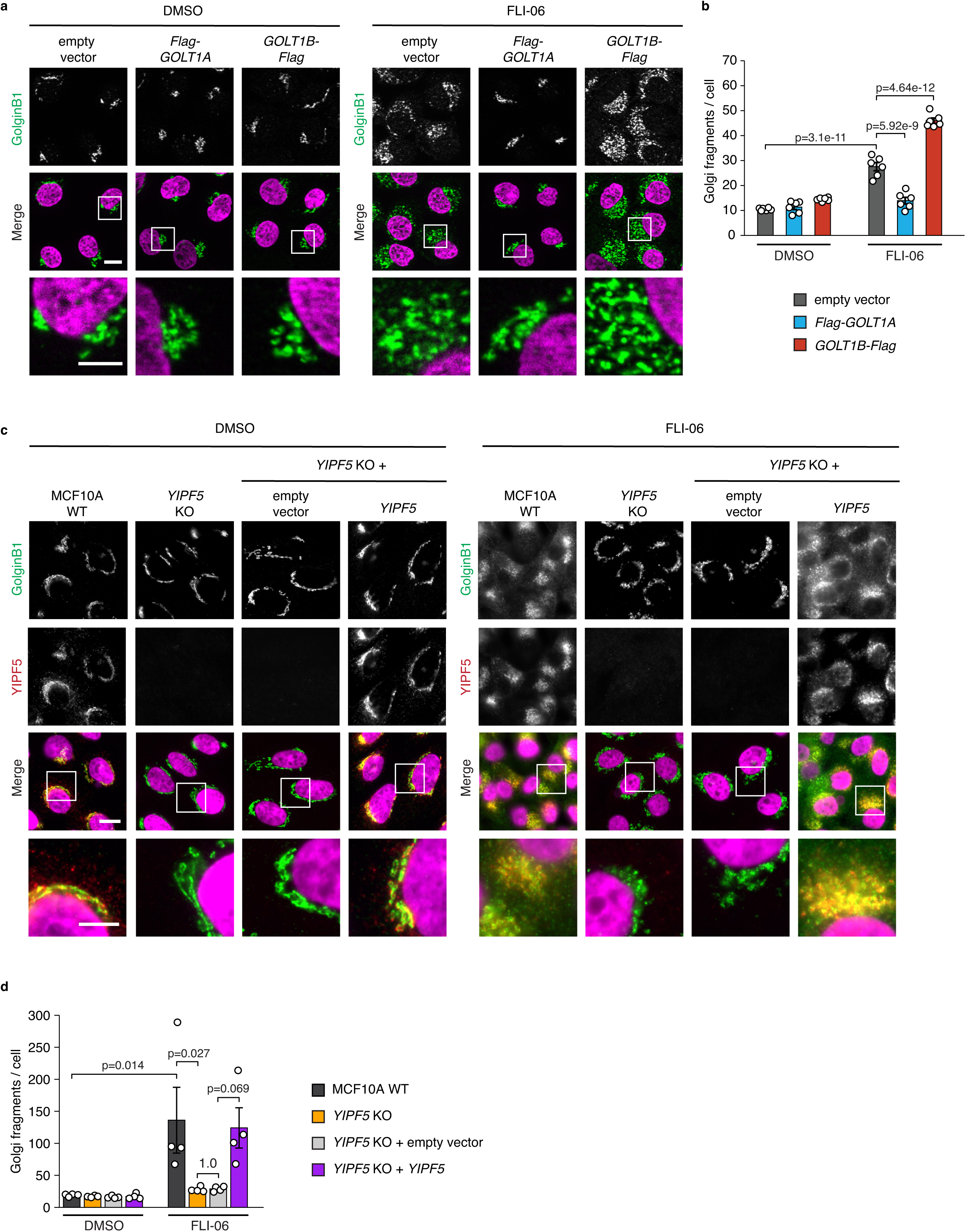
Overexpression of *GOLT1A* or knock-out of *YIPF5* rescue FLI-06 induced Golgi fragmentation. **a**) MCF10A cells stably expressing *Flag-GOLT1A*, *GOLT1B-Flag* or empty vector were treated for 24 h with 10 µM FLI-06 or DMSO as control, fixed and processed for immunofluorescence staining against the Golgi marker GolginB1 (green). Counterstaining with Hoechst-33342 (magenta) visualizes nuclei. Representative images of six biological replicates are shown. **b**) Quantification of Golgi-fragments based on immunofluorescence images of **a**). **c-d**) Immunofluorescence images (**c**) and quantification (**d**) of Golgi fragmentation of *YIPF5* KO and *YIPF5* or empty vector re-expressing MCF10A cells as described for **a**) and **b**). Data are shown as mean of six (**b**) or four (**d**) biological replicates and error bars represent SEM. Statistical testing: one-way-ANOVA test with Tukey HSD post-hoc test. Scale bar: 10 µm. Inserts (white square, scale bar: 5 µm) show higher magnification of the respective area.

**Fig. 3:**
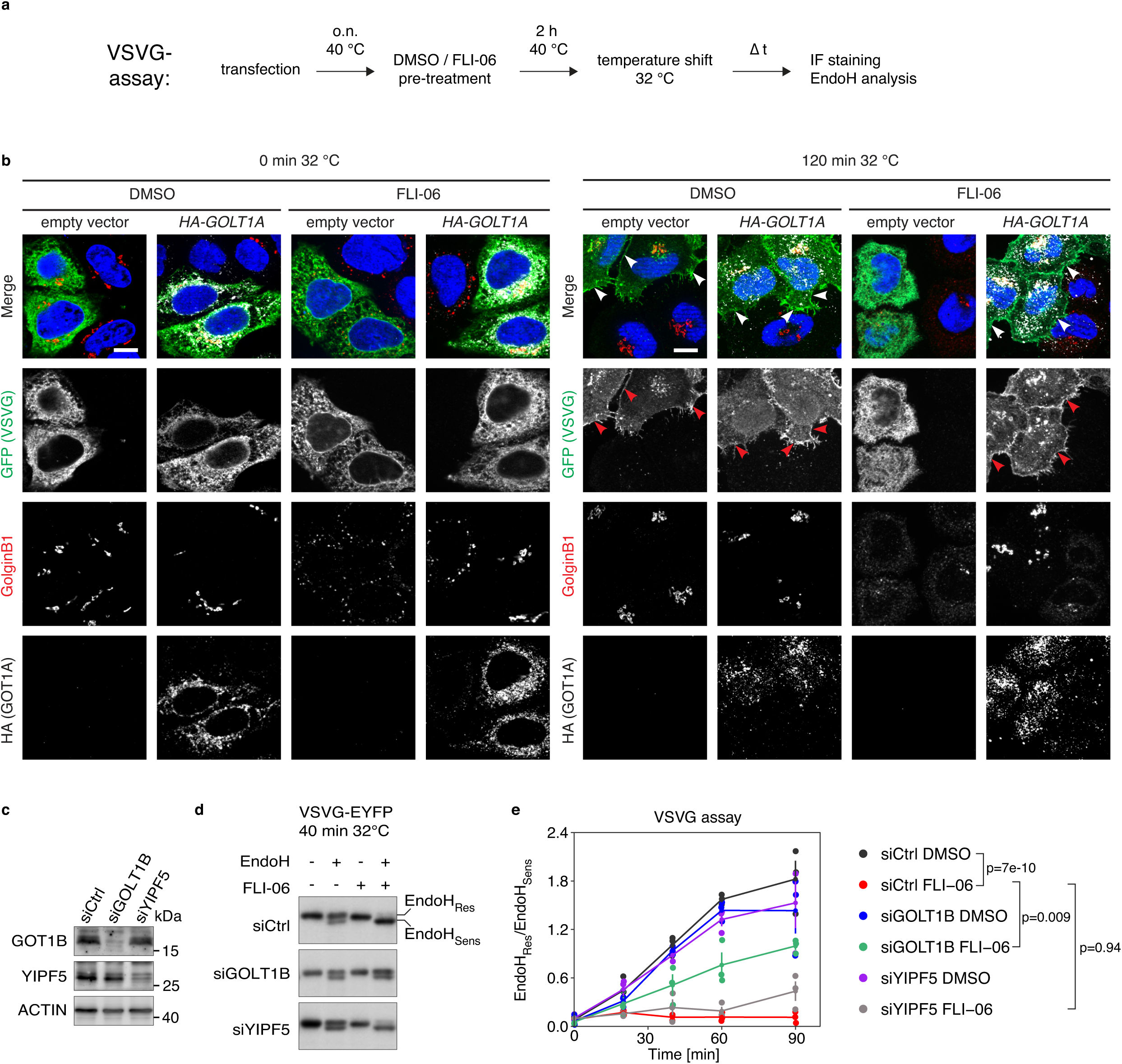
Overexpression of *GOLT1A* or knock-down of *GOLT1B* or *YIPF5* rescue FLI-06 induced ER-export block. **a**) Scheme of VSVG assay to monitor ER-export. **b**) HeLa cells were co-transfected with VSVG-EYFP and *HA-GOLT1A* or empty vector and incubated overnight (o.n.) at 40 °C. Cells were treated for an additional 2 h with 10 µM FLI-06 or DMSO at 40°C before the temperature shift to 32 °C for 2 h was performed. Thereafter cells were fixated and processed for immunofluorescence with antibodies against anti-HA (white), anti-GolginB1 (red) and anti-GFP (green). VSVG-EYFP localized to the plasma membrane is indicated by arrowheads. Counterstaining with Hoechst-33342 (blue) visualizes nuclei. Representative images of three biological replicates are shown. Scale bar: 10 µm. **c**) Immunoblotting analysis of siRNA knock-down of *GOLT1B* or *YIPF5* in HeLa cells. siCTRL, control siRNA. ACTIN, loading control. **d**) Immunoblotting analysis of EndoH resistant (Res) and sensitive (Sens) VSVG-EYFP in HeLa cells co-transfected with VSVG-EYFP and siRNAs as indicated. **e**) Quantification based on the immunoblots of **d**) at different time points after temperature shift to 32 °C. Data are shown as mean ratio of EndoH resistant and sensitive VSVG-EYFP of three biological replicates and error bars represent SEM. Statistical testing: one-way-ANOVA test with Tukey HSD post-hoc test.

Taken together, loss of YIPF5 or GOT1B expression and gain of GOT1A expression renders cells resistant to FLI-06 suggesting that these proteins are involved in a FLI-06-sensitive step in ER-export.

### GOT1A and GOT1B share a similar interactome and interact with each other

GOT1A and GOT1B are highly similar paralogues (Conchon et al., 1999) (Fig. S2A), yet have opposite functions in mediating FLI-06 resistance/sensitivity. To get more insights into the function of each, we first analyzed their subcellular localization in MCF10A cells stably overexpressing FLAG-GOT1A or GOT1B-FLAG (Fig. S3a,b, for functional validation of tagged constructs see Fig. S2h-k). Both colocalize to a similar degree with markers for the ER, ERES and Golgi, suggesting their different functions are not due to a different localization. We next were interested in determining the interactome of both proteins in the presence and absence of FLI-06. The stably overexpressing FLAG-GOT1A or GOT1B-FLAG MCF10A cells were subjected to immunoprecipitation with FLAG-antibody and subsequent mass-spectrometry analysis. FLAG-GOT1A interacted with 1337 proteins (Log_2_FC ≥ 0.58) in the absence and 1285 proteins (Log_2_FC ≥ 0.58) in the presence of FLI-06, with a high correlation coefficient of R^2^=0.88 of the two conditions (Fig. 4a, Fig. S4a,b Suppl. Table 1). Of the 27 known interaction partners of GOT1A (BioGRID) 5 were found in our co-immunoprecipitation. 86 of the 192 known GOT1B interaction partners (BioGRID), among them YIPF5, were also found to interact with GOT1A. As validation, we immunoprecipitated FLAG-GOT1A and blotted for YIPF5 and its interaction partner YIF1A (Yoshida et al., 2008). Both specifically co-precipitated with FLAG-GOT1A and did not change their interaction with GOT1A in the presence of FLI-06 (Fig. 4b).

**Fig. 4:**
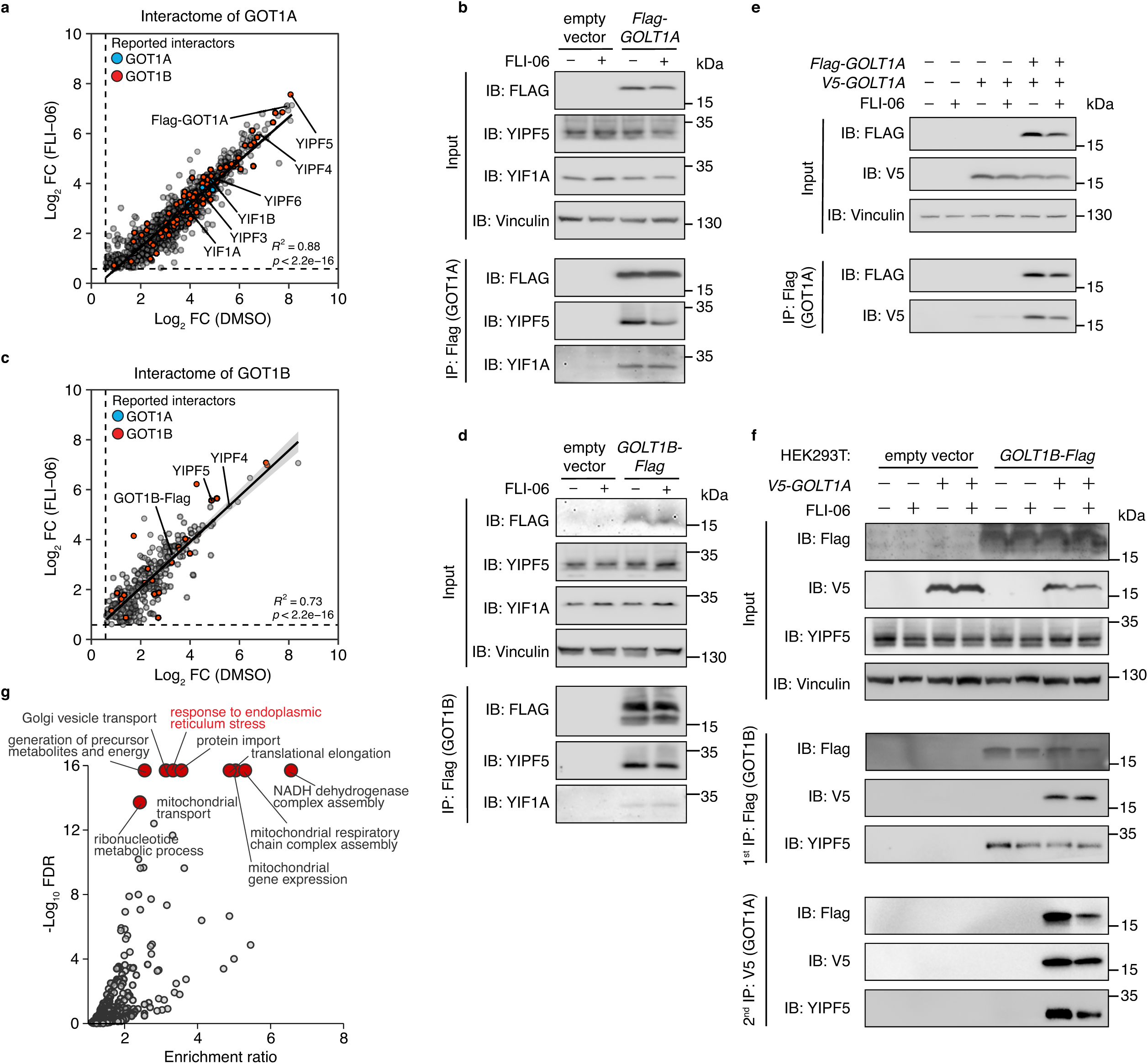
GOT1A and GOT1B have overlapping, yet distinct interactomes. **a**, **c**) Immunoprecipitation-mass spectrometry (IP-MS) analysis of MCF10A cells stably expressing *Flag-GOLT1A* (**a**) or *GOLT1B-Flag* (**c**). The cells were pretreated for 14 h with 10 µM FLI-06 or DMSO and IP was performed using Flag-antibodies. The Log_2_FC was calculated in comparison with empty vector expressing cells. Significant protein-protein interactors were defined as Log_2_ fold-change (FC) ≥ 0.58 and q-value ≤ 0.05. n = 4 biological replicates. Displayed are the Log_2_FC values of DMSO-against FLI-06-treated cells. Reported GOT1A (blue) or GOT1B (red) interactors (from BioGRID database) are highlighted. **b**, **d**) Co-IP of Flag-GOT1A (**b**) and GOT1B-Flag (**d**) in MCF10A cells as described for **a, c**) and immunoblotting analysis using the indicated antibodies. **e)** Co-IP and immunoblotting analysis of HEK293T cells co-transfected with *Flag-GOLT1A* and *V5-GOLT1A*. **f)** Consecutive Co-IP and immunoblotting analysis of HEK293T cells stably expressing *GOLT1B-Flag*, which were transfected with *V5-GOLT1A*. After IP against the Flag-tag (1^st^ IP), the eluate was used for an IP against the V5-tag (2^nd^ IP). All cells were pretreated for 14 h with 10 µM FLI-06 or DMSO before performing IP. Representative blots of three biological replicates are shown.

The interactome of GOT1B was overall smaller, with 320 proteins (Log_2_FC ≥ 0.58) in the absence and 317 proteins (Log_2_FC ≥ 0.58) in the presence of FLI-06, respectively (Fig. 4c, Fig. S4c,d, Suppl. Table 1). 175 (DMSO) or 222 (FLI-06 treated) proteins were found to interact both with GOT1A and GOT1B (Fig. S4 e,f) The correlation coefficient R^2^ was lower than in GOT1A, indicating a stronger change in the interactome by FLI-06. Like for GOT1A, also for GOT1B validation experiments demonstrated co-immunoprecipitation of endogenous YIPF5 and YIF1A, both known interaction partners (Hein et al., 2015) (Fig. 4d).

Interestingly, GOT1A interacted with itself and GOT1B, as indicated by co-immunoprecipitation experiments with co-expressed FLAG- and V5-tagged GOT1A (Fig. 4e) and GOT1B (Fig. 4f), respectively. The amount of co-immunoprecipitated YIPF5 bound to GOT1B is independent of the presence or absence of GOT1A in the complex, suggesting it interacts with both. The data also suggest that GOT1A overexpression replaces GOT1B in the complex, and vice versa. Interestingly, we noted that FLI-06 apparently destabilized the V5-GOT1A/GOT1B-FLAG/YIPF5 complex, as a second consecutive immunoprecipitation of the V5-GOT1A/GOT1B-FLAG/YIPF5 complex with V5 antibody precipitated less GOT1B-FLAG and YIPF5 when the complex was isolated from FLI-06 treated cells (Fig. 4f).

Surprisingly, gene-ontology analysis of the GOT1A interactome revealed an ER-stress signature (Fig. 4g) indicating the GOT1A/B YIPF5 complex could be involved in sensing and/or mediating ER stress signaling in response to FLI-06.

Taken together, the data suggest that GOT1A and GOT1B function in multi-subunit heteromeric complexes. The overlapping but not identical interactomes of GOT1A and GOT1B suggest that GOT1A/GOT1A or GOT1A/GOT1B or GOT1B/GOT1B-containing complexes perform different functions. The interactomes further suggest that GOT1A and GOT1B might interact with the ER stress machinery.

### RNAseq identifies a unique FLI-06 gene expression signature different from thapsigargin

The block of ER-export by FLI-06 is expected to result in a massive accumulation of newly synthesized proteins and a subsequent induction of ER stress. However, in previous studies we showed that all three branches of the unfolded protein response (UPR) are only mildly induced (Krämer et al., 2013). Yet, the interactomes of GOT1A/B indicated a connection to the ER stress machinery (Fig. 4g). To test whether GOT1A/B are involved in the regulation of the ER stress machinery and its target genes, we complemented the interactome studies by performing RNA-Seq analysis of MCF10A cells under ER stress. To this end, we treated MCF10A cells with DMSO, FLI-06 or Thapsigargin (TG), a known ER stress inducing agent (Fig. 5). Principle component analysis of all three conditions resulted in a clear separation of the conditions (Fig. 5a). FLI-06 induced the expression of 917 genes and reduced the expression of 469 genes (Fig. 5b), whereas TG resulted in larger changes (down: 1808/up: 2203 genes, Fig. 5c). Venn-diagram display of the data showed an overlap of 461/274 common up- or down-regulated genes, respectively (Fig. 5d). Pathway analysis of up-regulated genes revealed in both cases a strong relation to the ER, vesicular transport, ER stress, confirming TG as ER stress inducer and confirming that FLI-06 is a specific inhibitor with an ER-related target (Fig. 5g,h). Since both FLI-06- and TG-treated cells showed a clear ER stress signature, and since we showed earlier that FLI-06 only mildly induces stress we compared the effect size of those genes with a GO-term annotation “response to ER-stress” that were commonly up-regulated in FLI-06 and TG. Fig. 5i shows that on average FLI-06 induces ER stress genes less potently, confirming earlier findings (Krämer et al., 2013). Hierarchical clustering and gene ontology analysis confirms that FLI-06, just like TG induces expression of ER stress genes, but FLI-06 induces a specific gene expression signature that is different from TG and also tends to be weaker (Fig. 5f-i).

**Fig 5:**
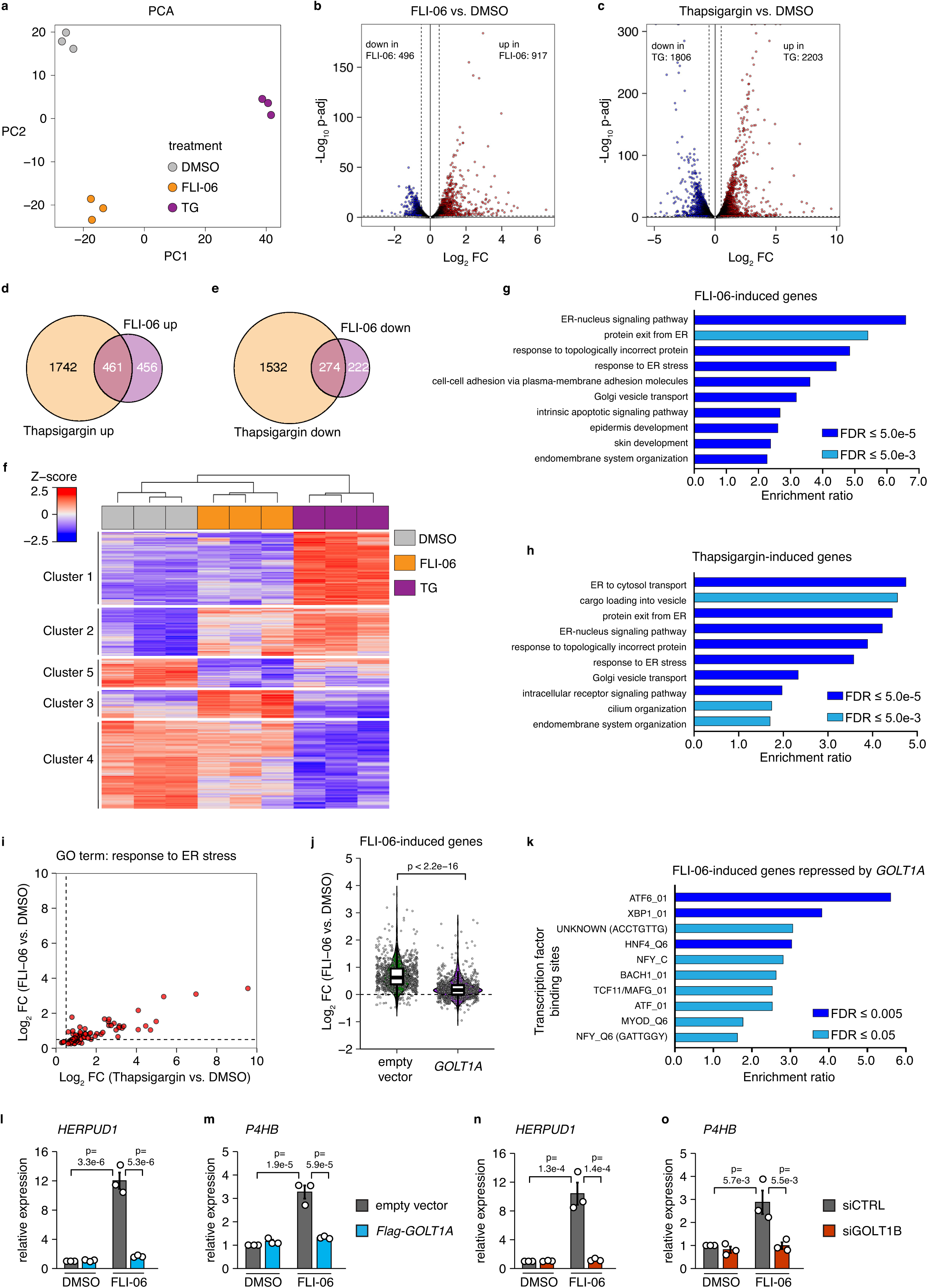
FLI-06 elicits a specific gene expression signature distinct from the ER-stress inducer Thapsigargin. **a**) PCA plot of RNA seq from MCF10A cells treated with FLI-06 (10 µM), Thapsigargin (100 nM) or DMSO (1:1000, control) for 6 h (n = 3 biological replicates). **b, c**) Volcano plot showing differentially expressed genes (p-adj < 0.01, Log_2_FC > 0.5 (up) / < −0.5 (down)) upon FLI-06 (**b**) or Thapsigargin treatment (**c**). Displayed is the Log_2_FC vs. −Log_10_ p-adj. **d, e**) Venn diagram showing the overlap of significantly upregulated (**d**) or downregulated (**e**) genes upon FLI-06 or Thapsigargin treatment, respectively. **f**) Heatmap of RNA seq data showing clusters of differentially expressed genes. **g, h**) GO term analysis for biological processes associated with FLI-06- (**g**) or Thapsigargin-induced genes (**h**). **i**) Scatterplot showing the differential expression of ER stress-associated genes (GO term: 0034976) upon FLI-06 or Thapsigargin treatment. Displayed is the Log_2_FC Thapsigargin vs. DMSO against Log_2_FC FLI-06 vs. DMSO. **j**) Violin plot of RNA seq from MCF10A cells stably expressing *Flag-GOLT1A* or empty vector treated with 10 µM FLI-06 for 6 h. The data are filtered for FLI-06-induced genes (defined in Fig. **5b**). Statistical testing: two-sample Wilcox test, n=3 biological replicates. **k**) From Fig. **5j** a gene set of FLI-06-induced genes, which were not anymore upregulated upon *GOLT1A* overexpression, was stratified. For this gene set, a GO term analysis for transcription factor binding sites in the promoters of genes was performed. **l-o**) qPCR expression analysis of *HERPUD1* (**l, n**) and *P4HB* (**m, o**) in MCF10A cells, which are stably expressing *Flag-GOLT1A* (**l, m**) or siGOLT1B (**n, o**) transfected. Cells were treated with FLI-06 (10 µM) for 14 h. Data are shown as mean of three biological replicates and error bars represent SEM. Statistical testing: one-way-ANOVA test with Tukey HSD post-hoc test.

We next tested how overexpression of *GOLT1A* affects genes induced by FLI-06, and we found that the gene signature induced by FLI-06 is to a large extent normalized by the overexpression of *GOLT1A* (Fig. 5j). *In silico* analysis of promoters of *GOLT1A*-repressed genes otherwise induced by FLI-06 revealed highly significant enrichment of ATF6 and XBP1 binding sites, two major transcriptional mediators of UPR (Hetz et al., 2020, Ron & Walter, 2007). Detailed qPCR analysis of *HERPUD1* and *P4HB* - two canonical ER stress genes upregulated in FLI-06 and TG-treated cells - showed that the attenuated induction after *GOLT1A* overexpression is much more pronounced in FLI-06 compared to TG-treated cells (Fig. 5l-m, S4g,h). Analogous results were obtained after *GOLT1B* knockdown, suggesting a similar mechanism (Fig. 5n,o, S4i,j). This suggests that GOT1A is not a general cell survival factor but rather a specific mediator of a non-canonical ER stress response after secretion blockade.

Taken together, the data suggest that FLI-06 elicits an ATF6 signature and that overexpression of *GOLT1A* or knock-down of *GOLT1B* suppressed ATF6 and/or XPB1 target genes after FLI-06 treatment.

### A transcription-independent role for ATF6 in FLI-06-mediated cell death

*GOLT1A* overexpression rescues FLI-06-mediated cell death and suppresses activation of ATF6 and/or XBP1 target genes. To independently validate ATF6 activation by FLI-06 and suppression by *GOLT1A*, we performed CUT&RUN experiments with antibodies against ATF6 and XBP1, respectively (Fig. 6a-c). In FLI-06-treated cells, ATF6 bound to 555 genomic sites and this number was strongly reduced (by ∼75%) to 136 peaks in *GOLT1A*-overexpressing cells (Fig. 6a,c). Consistent with stronger effects in our RNA-Seq analyses, TG treatment led to a higher number of peaks (n=1,331) which was also reduced by *GOLT1A* overexpression (n=735) but to weaker extent (by ∼45%). XBP1 binding, on the other hand, was not induced by FLI-06 but potently induced by TG treatment (Fig. 6b,c), demonstrating that the low number of XBP1 peaks in the FLI-06 condition is not simply due to low sensitivity of the antibody. GOT1A overexpression furthermore only marginally impacted on XBP1 binding in TG-treated cells (Fig. 6b,c). Consistent with our RNA-Seq data, overexpression of GOT1A was able to specifically blunt ATF6 recruitment to direct ATF6 targets induced by FLI-06 but failed to attenuate ATF6 binding after TG treatment (Fig. 6d).

**Fig. 6:**
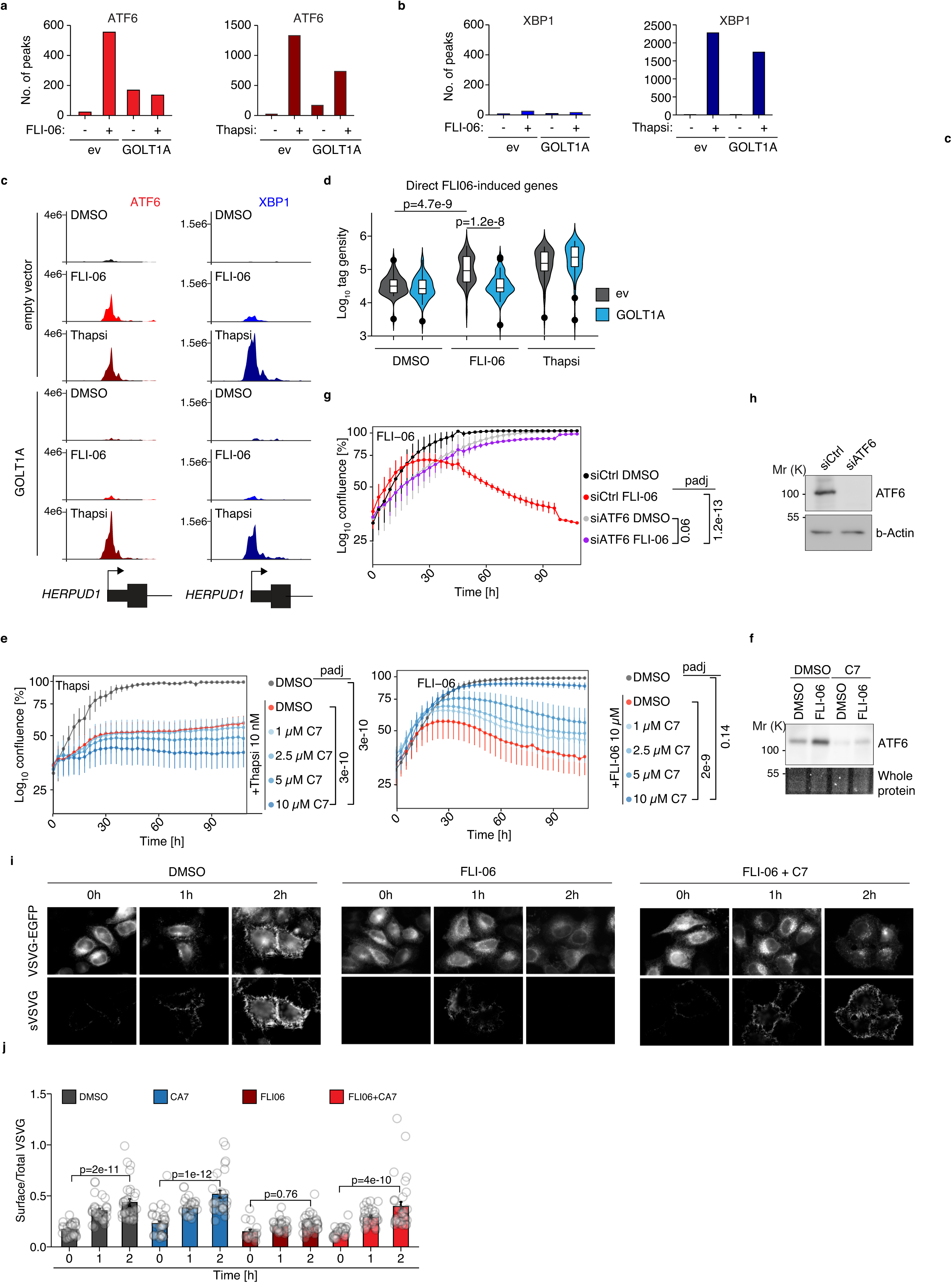
ATF6 is directly involved in the FLI-06 mediated secretion inhibition. **a,b)** Number of peaks of CUT&RUN experiments for ATF6 and XBP1 in MCF10A cells expressing empty vector (ev) or overexpressing GOLT1A in the presence or absence of 10 µM FLI06 or 100 nM Thapsigargin (Thapsi) for 3 hours. **c)** Representative CUT&RUN sequencing tracks at the FLI-06-induced gene *HERPUD1*. **d)** Normalized read counts in the ATF6 peaks close to promoters of FLI-06-induced genes. One-way ANOVA with post-hoc Tukey HSD. **e)** Incucyte growth curves of MCF10A cells treated with 10 nM Thapsigargin or 10μM FLI-06 in combination with DMSO or increasing concentrations of ATF6 inhibitor CeapinA7 (C7) as depicted. Plotted are the mean relative confluencies of three independent experiments. Error bars reflect SEM, two-way ANOVA. **f)** Immunoblot analysis of MCF10A cells treated with 10 µM FLI-06 ± 10 µM CeapinA7 (C7) for 6 hours. **g)** MCF10A cells transfected with control siRNA (siCtrl) or siRNA against ATF6 (siATF6) for 48h were re-plated, treated with 10 μM FLI-06 and proliferation was analyzed using IncuCyte microscopy for another 100h. Plotted are the mean relative confluencies of three independent experiments. Error bars reflect SEM, two-way ANOVA. **h)** Western Blot of cell lysates of control siRNA or ATF6-siRNA transfected MCF10A cells using antibodies against ATF6 or actin as loading control. **i)** VSVG-EYFP transport assay of Hela cells transiently transfected with VSVG-EYFP, incubated at 40°C overnight, treated 30 min before shifting to the permissive temperature with 40 μg/ml cycloheximide and DMSO or 10μM FLI-06 in combination with 10μM CeapinA7 (C7). Cells were then chased in the presence of indicated drugs for 1h or 2h at 32°C. Surface VSVG was stained by anti-VSVG-antibody EB0010 before fixation and cells processed for immunofluorescence. **j)** Depicted is the mean ratio of total VSVG-EYFP and surface-localized VSVG-EYFP from n=3 independent experiments from i). Open circles reflect regions of interest, each containing 5-140 transfected cells. Two-way ANOVA test with post-hoc Tukey HSD.

ATF6 is a membrane-bound, ER-localized transcription factor which is recruited to ERES upon activation, translocates to the Golgi where it is cleaved by intramembrane proteases S1P and S2P (Haze et al., 1999, Ye et al., 2000). The cleavage releases the N-terminal, transcriptionally active domain which is then transported into the nucleus (Haze et al., 1999). To test whether ATF6 activation is involved in FLI-06-mediated cell death, we inhibited ATF6 translocation and processing with CeapinA7 (Gallagher & Walter, 2016) and assayed cell survival in the presence of FLI-06 or TG, respectively. As expected, simultaneous treatment of cells with TG and CeapinA7 reduced their viability in a dose-dependent manner compared with TG alone (Fig. 6e). In contrast and surprisingly, cells incubated with FLI-06 were rescued when ATF6 signaling was inhibited (Fig. 6e), and FLI-06 treatment induced ATF6 protein which was attenuated in C7-treated cells (Fig. 6f). Knockdown of ATF6 also rendered cells resistant to FLI-06, confirming specificity of CeapinA7 (Fig. 6g,h).

### ATF6 acts as an ER protein, not a transcription factor, in FLI-06 mediated ER-export block

We next tested whether inhibition of ATF6 would overcome the ER-export block by FLI-06 or would rescue the cells via different mechanisms. To this end, we used the VSVG-EGFP transport assay in the presence of CeapinA7 and cycloheximide (CHX) to inhibit translation. The efficacy of CHX was shown by the complete loss of the short-lived protein c-MYC (Fig. S5a). Notably, CeapinA7 in combination with FLI-06 partially rescued the ER-export inhibition of VSVG-EGFP (Fig. 6i, j). Moreover, CeapinA7 was added only 30min before transferring cells to permissive 32°C and that was sufficient to release VSVG-EGFP in the presence of FLI-06, excluding it is based on ATF6-downstream transcription. Also, the presence of CHX during the 32°C prevented a transcriptional/translational response downstream of ATF6.

We noticed that FLI-06-treated cells had higher ATF6 levels (∼2-fold after 3 h of FLI-06-treatment, Fig. 6f). Increasing ATF6 expression alone, however, is not sufficient to block ER-export as shown by overexpressing GFP-ATF6 in a cell line expressing secreted alkaline phosphatase (SEAP) as a marker for secretion (Fig. S5b,c) and by co-expressing VSVG-mCherry and GFP-ATF6 (Fig. S5d). This suggests that the increase in ATF6 levels by FLI-06 is essential but not sufficient for the ER export block. Taken together, our data suggest that GOT1A and GOT1B link ER export to membrane-bound ATF6, demonstrating a hitherto undescribed role for ATF6 in ER-export, independent of its well-established role as a UPR-transcription factor.

## Discussion

Chemical genetic screens using CRISPR/Cas technology have been widely used for target identification (reviewed in Jost & Weissman, 2018). We applied the principle to find putative targets of FLI-06, an inhibitor of cargo recruitment into ERES (Krämer et al., 2013, Yonemura et al., 2016). The most significantly enriched genes are encoding the small polytopic membrane proteins GOT1A, GOT1B and YIPF5. All three are involved in a critical step in FLI-06 mediated ER-export inhibition, since knockout or knockdown of *GOLT1B* or *YIPF5* or overexpression of *GOLT1A* rendered cells resistant to FLI-06.

GOT1B and YIPF5 localize to the early secretory pathway and interact with inner COPII coat components SEC23/24 (Tang et al., 2001, Wang et al., 2016). They interact genetically (Lorente-Rodriguez et al., 2009) and directly (Hein et al., 2015) with each other but their precise role has not been defined. Our results show that GOT1A, GOT1B and YIPF5 localize to the ER, the ERGIC and the early Golgi, and YIPF5 additionally is also enriched in ERES. All three interact with each other, as shown before for YIPF5 and GOT1B (Hein et al., 2015). Knock-down of *GOLT1B* or knock-out of YIPF5 has no effect on proliferation and VSVG-EYFP transport out of the ER, confirming earlier findings (Kranjc et al., 2017, Yoshida et al., 2008) and suggesting they are not part of the core ER-export machinery.

We concentrated in this study on GOT1A/B because of their high homology but opposing role in our assay. Not being part of the core ER export machinery, GOT1A/B might be molecular mediators of the known link between secretion and ER stress (Shaheen, 2018). Perturbed ER-export, in our case induced by FLI-06, causes the accumulation of properly folded, export-competent proteins in the ER. This is different from the accumulation of misfolded proteins caused by mutations in cargo proteins or by changes in the ER milieu. Indeed, our data show that blocking ER-export by FLI-06 causes a distinct ER stress signature, different from Thapsigargin (TG). TG is a classical ER stress inducer that dramatically increases unfolded proteins (Foufelle & Fromenty, 2016). As shown before (Krämer et al., 2013), FLI-06 activates ER stress to a lesser extent than TG, likely because the classical activation of UPR is based on the activation of ATF6, IRE1 and PERK via sensing unfolded proteins (Hetz et al., 2020, Ron & Walter, 2007). FLI-06 does not inhibit protein folding in the ER (Yonemura et al., 2016), therefore most likely folded proteins accumulate in the ER. Fully folded, export-competent proteins will not bind BIP and therefore they will only mildly activate the UPR sensors, possibly explaining why the FLI-06 gene expression signature is weaker than the one induced by Thapsigargin.

Overexpression of GOLT1A or knockdown of GOLT1B also reduced the expression of TG-induced HERPUD1 and P4HB, but to a much lesser extent than in the case of FLI-06 treatment. This suggests a role for GOLT1A/B also in the classical unfolded protein response that requires further elucidation.

Interestingly, the ATF6 ER-stress signature we observed after treatment with FLI-06 normally involves translocation of ATF6 from the ER to the Golgi, but that transport step is blocked by FLI-06. ATF6 signaling could be activated by some ATF6 molecules escaping the FLI-06 block and reaching the Golgi. Alternatively, under these special blocking conditions ATF6 can be activated already in the ER, maybe because also its proteases, S1P and S2P (Ye et al., 2000), also become trapped in the ER. Such a cleavage in the ER was observed when S1P and S2P were re-localized to the ER by BFA (Gallagher & Walter, 2016).

Surprisingly, we found that knocking down ATF6 or inhibiting it with CeapinA7 rescued the FLI-06-dependent ER-export blockage. Moreover, our data strongly suggest that CeapinsA7’s effect is ER-based and independent of ATF6’s role as a transcription factor. This is supported by the fact that a short incubation time of only 30min of CeapinA7 before the temperature shift from 40° to 32°C resulted in a rescue of ER-export of VSVG-EGFP, even in the absence of any protein translation. Strikingly, ATF6 levels were strongly increased after FLI-06 treatment, indicating a stabilization, maybe by an interacting protein. Apparently, the increased levels of ATF6 are not sufficient to block ER-export since overexpression of ATF6 alone did not block secretion. Since knockdown or inhibition of ATF6 rescues FLI-06 secretion inhibition, the accumulation of ATF6 likely is necessary but not sufficient for the ER-exit block. This non-canonical function of ATF6 needs to be further explored in future studies.

What is the molecular function of GOT1A and GOT1B, if they are not part of the basic ER-export machinery? They could be auxiliary proteins or cargo receptors with a very specific set of cargoes not essential in cell lines. Or they could be part of a sensor measuring the rate of ER-export and fine-tune the expression or availability of proteins needed for export (Subramanian et al., 2019). Quality control in the ER is critical for cellular function (Ellgaard & Helenius, 2003) and the presence of an additional control system has been experimentally demonstrated, but not fully elucidated (Mezzacasa & Helenius, 2002). The GOT1A/B/YIPF5 complex could be part of this mechanism. FLI-06 could directly bind GOT1B and/or YIPF5, causing a clogging of ERES and thereby preventing export. Alternatively, if the GOT1A/B/YIPF5 complex is part of a feedback mechanism, overactivation of such a mechanism by FLI-06 could cause a complete shut-down of ER-export. Knockdown of *GOLT1B* could simply remove the target for FLI-06 or disrupt the feedback mechanism. Transport would resume, and since the ER-export sensor is not essential, cells will be resistant to FLI-06 and survive. GOT1A could have the same function like GOT1B, but since it is highly homologous but not identical, it could lack the binding site for FLI-06 or the binding site for a FLI-06 target protein. Since GOT1A and GOT1B bind to the same complex containing YIPF5 (Fig. 4), overexpression of GOT1A could replace GOT1B, rendering cells resistant to FLI-06. Future work, for example using cross-linkable FLI-06 derivatives, will help to identify the direct target. Previously the ryanodine receptors were proposed as targets for FLI-06, but their role in ER export and/or cell survival was not elucidated (Gunaratne et al., 2022). Also we found no evidence for a role of ryanodine receptors in ER export. In sum, we identify here GOT1A, GOT1B and YIPF5 as non-essential components of the early secretory pathway that are involved in the ER-exit export block caused by FLI-06. In addition, we propose a novel function of full-length ATF6 in ER-export, independent of its well characterized role as sensor for unfolded proteins and transcription factor for UPR-genes.

## Materials and methods

### Antibodies, primers and plasmids used

See supplementary experimental procedures.

### Cloning

Techniques of restriction cloning were used throughout. sgRNAs for SAM-mediated endogenous gene overexpression were cloned into the lenti_sgRNA(MS2)_zeo_backbone (kind gift from Feng Zhang, Addgene #61427) as reported previously (Joung et al., 2017). Sequences of SAM sgRNAs were obtained from the SAM library (Konermann et al., 2016) and SAM non-targeting sgRNAs (sgNT) were previously published (Joung et al., 2017). For shRNA expression, the lentiviral SGEP vector (kind gift from Johannes Zuber, Addgene #111170) was used. For exogenous gene overexpression, the coding sequence of genes was PCR-amplified (Phusion™ Hot Start II DNA Polymerase, Thermo Fisher Scientific, cat.no. F549L) from human cDNA and cloned into the LeGO-iG2 (kind gift from Boris Fehse, Addgene #27341). YIPF5, GOT1A or GOT1B were amplified from cDNA using appropriate primers. To add epitope tags to YIPF5, GOT1A or GOT1B, the tag sequence was included in PCR primers used for amplification of the cDNA. Plasmid DNA was extracted using the QIAGEN Plasmid Maxi Kit (Qiagen, cat.no. 12165). All cloned sequences were validated by Sanger sequencing.

### Mammalian Tissue Culture and transfection

All cell lines were cultivated at 5 % CO_2_, 95 % relative humidity and 37 °C in a cell culture incubator. Mammalian cell lines were handled under sterile conditions using a laminar flow hood and the cells were regularly tested for mycoplasma contamination with a PCR-based mycoplasma detection kit (Minerva-Biolabs, cat.no. 11-1250). Subcultivation was performed every 2-4 days at a confluence of maximum 80 % using trypsin (Sigma-Aldrich, cat.no. 59427C-100ML). HEK293T-LentiX (Takara Bio, cat.no. 632180) and HeLa (Unauthenticated HeLa-Kyoto, kind gift from Rainer Pepperkok, EMBL Heidelberg, Germany) cells were cultivated in DMEM (high glucose, GlutaMax, Thermo Fisher Scientific, cat.no. 61965059) supplemented with 10 % FBS (Thermo Fisher Scientific, cat.no. 10270106) and 1 % penicillin-streptomycin (Thermo Fisher Scientific, cat.no. 15140122). MCF10A cells (kind gift from Almut Schulze (University of Würzburg, Germany), authenticated using STR profiling) were cultivated in DMEM supplemented with 5 % horse serum (Thermo Fisher Scientific, cat.no. 16050122), 1% penicillin-streptomycin, 10 µg/ml human insulin (Sigma-Aldrich, cat.no. I9278), 0.5 µg/ml hydrocortisone (Sigma-Aldrich, cat.no. H0888-5G), 0.1 µg/ml cholera toxin (Sigma-Aldrich, cat.no. C8052-2MG) and 20 ng/ml human EGF (Biomol, cat.no. 50349.1000). BJ cells (ATCC cat.no. CRL-2522) were cultivated in DMEM (High Glucose, Sigma Aldrich, cat.no. D5648) supplemented with 10% FBS and 1 % penicillin-streptomycin.

Reverse transfection of cells with ON TARGETplus siRNAs (Dharmacon, final concentration: 10 nM) was performed for 12 h with Lipofectamine™ RNAiMAX (Thermo Fisher Scientific, cat.no. 13778030) as described by the manufacturer. Transient transfection of cells with plasmid DNA was performed on 80 % confluent cells using PEI-Max (Polyscience, cat.no. 24765-1, DNA:PEI ratio of 1:3) for 12 h. For the time of transfection, the cells were kept in transfection medium. The transfection medium of HEK293T-LentiX and HeLa cells consisted of DMEM supplemented with 2% FBS and without penicillin-streptomycin. For MCF10A cells, the transfection medium contained the same components as the culture medium but lacked penicillin-streptomycin and cholera toxin.

### Lentiviral transduction

HEK293T-LentiX cells were co-transfected for 12 h with 10 µg psPAX2 (kind gift from Didier Trono, Addgene #12260), 2.5 µg pMD2.G (kind gift from Didier Trono, Addgene #12259) and 10 µg lentiviral plasmid using PEI-Max (Polyscience, cat.no. 24765-1). Lentiviral supernatant was harvested 36, 48 and 60 h after transfection. After pooling and sterile-filtering the supernatant, the cells were infected for 24 h with viral supernatant, which was diluted in culture medium supplemented with 8 µg/ml of protamine sulfate (Sigma-Aldrich, cat.no. P4505-1G). To generate stable cell lines, selection with the appropriate selective antibiotics was performed.

### SAM Screening

The genome-wide human SAM sgRNA library (kind gift from Feng Zhang, Addgene #1000000057) was amplified as previously described (Joung et al., 2017). A balanced library distribution was verified by NGS. Clonal MCF10A cell lines (MCF10A-SAM #9 and #28) with stable lentiviral expression of the SAM components, dCas9-VP64 and MCP-p65-HSF1 (kind gift from Feng Zhang, Addgene #61425 and 61426), were infected with the lentiviral SAM library at a low viral titer (MOI=0.5) with a library coverage of at least 500-fold. For the screening, cells (>3.5 x 10^7^ cells to guarantee a 500-fold library coverage) were cultivated at a density of 1.5 x 10^4^ cells / cm^2^. Selection with 100 nM BFA (Sigma-Aldrich, cat.no. B7651-5MG) or 10 µM FLI-06 (Sigma-Aldrich, cat.no. SML0975-25MG) was started one day after plating and continued until clonal expansion of surviving cells was observed. Medium was changed every 48h. gDNA from cells selected with BFA or FLI-06 and unselected control cells were extracted using the QIAamp DNA Blood Maxi Kit (Qiagen, cat.no. 51194). The integrated sgRNA cassette was amplified in a first PCR. After pooling the product of the first PCR, a second PCR was performed to add Illumina adapters and barcodes for NGS. Quality check of libraries was done using an Agilent 2100 Bioanalyzer Instrument and a DNA 7500 assay (Agilent Technologies). Quantification was done using a RT-qPCR assay. NGS was conducted on a NextSeq^®^ 500 (Illumina, SY-415-1001) using the High Output Kit v2.5 (Illumina, cat.no. 20024906) with 75 cycles of single-end sequencing. Sequence information was converted to FASTQ format using bcl2FastQ v2.19.1.403. Quality-filtering (>Q30) of the sequencing data and adapter removal was performed with cutadapt (v2.7). The filtered reads were mapped to a custom reference containing all sgRNA sequences in the SAM library using bowtie2. To all samples a pseudocount of +1 was added to avoid division by 0. The reads were normalized by sequencing depth (reads per sgRNA/ million mapped reads) and subsequently used in RSA analysis (Konig et al., 2007). For the RSA analysis, the following parameters were used: --l= median plus 1xSD, --u= median plus 3xSD.

### CRISPR-Cas9 knockout screen

The genome-wide human GeCKO lentiviral sgRNA library v2 (kind gift from Feng Zhang, Addgene #1000000048), consisting of the 2 pools A and B, was amplified as previously described (Joung et al., 2017) and the balanced library distribution was verified by NGS for both of them. The BJ cells were infected with the pseudoviral particles at low titer (MOI= 0.4), with a library coverage of at least 500-fold. For the screening the BJ cells (>3.5 x 10^7^ cells per pool to guarantee a 500-fold library coverage) were cultivated at a density of 9 x 10^4^ cells / cm^2^. Selection with Puromycin (5µg/ml) started 2 days after transduction and was kept until the end of the experiment. Immediately after selection 10 µM FLI-06 was applied; medium was changed every 48h. Selection continued until expansion of surviving cells was enough for the gDNA isolation. Cells transduced with the library Pool A didn’t survive to that extent. gDNA from cells selected with FLI-06 and unselected control cells were extracted using classical ethanol precipitation protocol. The integrated sgRNA cassette was amplified in a first PCR. After pooling the product of the first PCR, a second PCR was performed to add Illumina adapters and barcodes for NGS (Shalem et al., 2014). NGS was conducted on a MiSeq^®^. Quality check of libraries was done using an Agilent 2100 Bioanalyzer Instrument and a DNA 7500 assay (Agilent Technologies). Quantification was done using a RT-qPCR assay. Libraries were sequenced on a MiSeq System (Illumina) using a 150 cycle kit v3 (R1: 131; I1: 8 cycle) by spike-in 15 % Illumina’s PhiX library in order to improve the per cycle base composition. Sequence information was converted to FASTQ format using bcl2fastq v2.16. RSA analysis was performed identical to the SAM screen using RSA.

### Generation of CRISPR-Cas9 knockout cells

For generation of a vector for generating YIPF5 KO cells gRNA-coding oligonucleotides YIPF5 g57 BbsI Fwd/Rev or YIPF5 g60 BbsI Fwd/Rev were annealed and cloned in BbsI restriction enzyme cut pSpCas9(BB)-2A-GFP plasmid. MCF10A cells were transfected with the plasmids via nucleofection and the medium changed after 5h. Using the encoded GFP, cells were FACS-sorted 48h after transfection into 96-well plates with one cell/well and incubated with conditioned media from 40-50% confluent cells. After transferring to 6-well plates cells were cultivated in regular medium. DNA from single cell clones was genotyped by PCR using primers YIPF5 exon 3 Fwd/Rev. One clone from the g60-gRNA transfected cells was used for further analysis. This clone has a −20nt deletion in one allele and a +1 insertion in the other allele.

### RNA Expression analysis

Total RNA was extracted from cells with peqGOLD TriFast (VWR, cat.no. 30-2010). Synthesis of cDNA was performed with M-MLV Reverse Transcriptase (Promega, cat.no. M1705) and random hexamer primers (Sigma-Aldrich, cat.no. 11034731001-2MG). Gene expression was determined by qPCR using InnuMIX qPCR DSGreen Standard (Analytik Jena, cat.no. 845-AS-1320500) supplemented with ROX Reference Dye (Thermo Fisher Scientific, cat.no. 12223012) at a dilution of 1:50. The PCR and data acquisition were performed with a StepOnePlus™ Real-Time PCR System (Applied Biosystems, cat.no. 4376600). For each biological replicate of a given experiment, three technical replicates were performed. Beta-2-Microglobulin (B2M) was used as a reference gene for normalization. The relative expression was calculated by applying the ΔΔCT-method. Statistical analysis and visualization were performed with RStudio using in-house scripts.

### Immunoprecipitation

Cells were lysed with a buffer consisting of 50 mM Tris-HCL (pH 7.5), 150 mM NaCl, 1 mM MgCl_2_, 5 % (v/v) Glycerol, 1 % (v/v) NP-40, 1:1000 Protease inhibitor cocktail (Sigma-Aldrich, cat.no. P8340-5ML) and 1:1250 benzonase (Sigma-Aldrich, cat.no. E1014-25KU). After measuring the protein concentration with Pierce™ 660 nm Protein Assay Reagent (Thermo Fisher Scientific, cat.no. 22660), the concentration of the different samples was adjusted. Immunoprecipitation of Flag-tagged proteins was performed with Anti-Flag M2 Affinity Gel (Sigma-Aldrich, cat.no. A2220-1ML) at 4 °C with overhead rotation for 6 h. Then, unbound proteins were washed out 3x with 500 µl of lysis buffer. To elute precipitated proteins, the Affinity Gel was incubated with 2x vol of 100 mM glycine buffer (pH 2.5) at RT and 350 rpm agitation for 5 min. After centrifugation, the supernatant was collected with a Hamilton syringe and the elution repeated once. To neutralize the solution, 1 M unbuffered Tris was added to the eluate in a ratio of 1:5.

For consecutive Co-IP (Re-IP), the first IP against the Flag-tag was performed as described above with the following changes: Incubation of the protein lysate with the Anti-Flag M2 Affinity Gel was performed overnight (14 h). Then, the Anti-Flag M2 Affinity Gel was washed 3x with lysis buffer and 1x with TBS buffer. To elute precipitated proteins, the Anti-Flag M2 Affinity Gel was incubated with 2x vol of 100 µg/ml Flag-Peptide (in TBS, Sigma-Aldrich, cat.no. F3290) at 4 °C and 350 rpm agitation for 30 min. The elution was repeated once. To perform a secondary IP against the V5-tag, the eluate of the first IP was incubated with Anti-V5 Agarose Affinity Gel (Sigma-Aldrich, cat.no. A7345) at 4 °C with overhead rotation for 4 h. After 3x washing with 500 µl TBS, the proteins were eluted twice by incubation with 2x vol of 100 mM glycine buffer (pH 2.5) at RT and 350 rpm agitation for 5 min. To neutralize the solution, 1 M unbuffered Tris was added to the eluate in a ratio of 1:5.

### SDS-PAGE and Immunoblotting

Protein samples were diluted with 6x sample buffer (12 % SDS, 0.0004 % Bromphenol blue, 47% glycerol, 12 % Tris-HCl (pH 6.8), 9.3 % DTT), denaturated at 95 °C for 5 min and separated on Bis-Tris polyacrylamide gels using the Mini-PROTEAN^®^ Tetra Handcast System (Biorad, cat.no. 1658029FC). For visualization of total protein load, stain-free gel imaging was used. To this end, the gel was incubated in 10 % (v/v) trichloroacetic acid for 5 min and washed thoroughly with ddH_2_O. Fluorescence was induced and images were recorded using the GelDoc XR+ system (Biorad). From the gel, the proteins were transferred to 0.45 µm PVDF membranes (Merck Millipore, cat.no. IPVH00010). After blocking unspecific binding sites of the membrane at RT for 1 h with 5 % skim milk powder in TBS-T, the membrane was incubated overnight at 4 °C with primary antibodies diluted in 5 % BSA in TBS-T. Then, the membrane was washed 3x with TBS-T and probed with the appropriate horseradish peroxidase-coupled secondary antibodies diluted in 5 % skim milk TBS-T. The secondary antibody was incubated at RT for 1 h and unbound antibody was removed by 3x washing with TBS-T. Chemiluminescence was induced by incubation with Immobilon Western HRP Substrate (Merck Millipore, cat.no. WBKLS0500) and visualized with the myECL™ Imager (Thermo Fisher Scientific).

#### Proteomics

##### Sample preparation for proteome analysis

IP samples were mixed with 10x lysis buffer (final concentration: 100 mM HEPES, 50 mM DTT, 1 % (w/v) SDS) and snap frozen until processing. On preparation for MS, samples were thawed and sonicated (Bioruptor^®^ Plus, Diagenode) for 10 cycles (30 s ON/60 s OFF) at high setting, at 4 °C, followed by boiling at 95 °C for 5 min. Reduction was followed by alkylation with 15 mM iodoacetamide (IAA) for 30 min at room temperature in the dark. Proteins were precipitated overnight at −20 °C after addition of 5x volume of ice-cold acetone. The following day, the samples were centrifuged at 16’000 rcf and 4 °C for 30 min and the supernatant carefully removed. Pellets were washed twice with 300 µl of ice-cold 80 % (v/v) acetone in water then centrifuged at 16’000 rcf and 4 °C for 10 min. They were then allowed to air-dry before addition of 25 µl of digestion buffer (3 M Urea, 100 mM HEPES, pH 8.0). Samples were resuspended with sonication (as described above), LysC (Wako) was added at 1:100 (w/w) enzyme/protein ratio and digestion proceeded at 37 °C for 4 h with shaking (1000 rpm for 1 h, then 650 rpm). Samples were then diluted 1:1 with Milli-Q water and trypsin (Promega) added at the same enzyme/protein ratio. Samples were further digested overnight at 37 °C with shaking (650 rpm). The following day, digests were acidified by the addition of TFA to a final concentration of 10 % (v/v) and then desalted with Waters Oasis^®^ HLB µElution Plate 30 µm (Waters Corporation) in the presence of a slow vacuum. In this process, the columns were conditioned with 3x 100 µl of solvent B (80% (v/v) acetonitrile, 0.05 % (v/v) formic acid) and equilibrated with 3x 100 µl of solvent A (0.05 % (v/v) formic acid in Milli-Q water). The samples were loaded, washed 3 times with 100 µl of solvent A, and then eluted into 0.2 ml PCR tubes with 50 µl of solvent B. The eluates were dried down with the speed vacuum centrifuge and dissolved at a concentration of 1 µg/µl in reconstitution buffer (5 % (v/v) acetonitrile, 0.1 % (v/v) formic acid in Milli-Q water). Protein amount was estimated based on cell number input and it was confirmed with an SDS-PAGE gel of 4 % of each sample against an in-house cell lysate of known quantity. Reconstituted peptides were analyzed by data independent acquisition (DIA).

##### Data-independent acquisition and data processing

Reconstituted peptides were spiked with retention time iRT kit (Biognosys AG), and separated using the nanoAcquity UPLC system (Waters) with a trapping (nanoAcquity Symmetry C18, 5 µm, 180 µm x 20 mm) and an analytical column (nanoAcquity BEH C18, 1.7 µm, 75 µm x 250 mm). The outlet of the column was coupled to a Q exactive HF-X (Thermo Fisher Scientific) using the Proxeon nanospray source. Samples were loaded at constant flow of solvent A (0.1 % (v/v) formic acid in Milli-Q water) at 5 µl/min onto the trap for 6 min. Peptides were eluted via the analytical column at 0.3 µl/min and introduced via a Pico-Tip Emitter 360 µm OD x 20 µm ID; 10 µm tip (New Objective). A spray voltage of 2.2 kV was used. During the elution step, the percentage of solvent B (acetonitrile with 0.1 % (v/v) formic acid) increased in a non-linear fashion from 0 % to 40 % in 60 min. Total run time was 75 min. The capillary temperature was set at 300 °C.

DIA acquisition was run with the following MS conditions. The spray voltage was 2.2 kV und the RF ion funnel was set to 40 %. Full scan MS spectra with mass range 350-1650 m/z were acquired in profile mode in the Orbitrap with resolution of 120000 FWHM. The filling time was set at maximum of 60 ms with limitation of 3.0 x 10^6^ ions. DIA scans were acquired with 30 mass window segments of differing widths across the MS1 mass range. The default charge state was set to 3+. HCD fragmentation (stepped normalized collision energy; 25.5, 27, 30 %) was applied and MS/MS spectra were acquired with a resolution of 30000 FWHM with a fixed first mass of 200 m/z after accumulation of 3.0 x 10^6^ ions or after filling time of 47 ms (whichever occurred first). Data were acquired in profile mode. For data acquisition and processing of the raw data Xcalibur 4.0 (Thermo Fisher Scientific) and Tune version 2.9 were employed.

Acquired data were processed using Spectronaut Professional v13.10 (Biognosys AG). Raw files were searched by directDIA search with Pulsar (Biognosys AG) against the human UniProt database (Homo sapiens, entry only, 20.186 entries, release 2016_01) with a list of common contaminants appended, using default settings. For quantification, default BGS factory settings were used, except: Proteotypicity Filter = Only Protein Group Specific; Major Group Quantity = Median peptide quantity; Major Group Top N = OFF; Minor Group Quantity = Median precursor quantity; Minor Group Top N = OFF; Data Filtering = Qvalue percentile with Fraction = 0.2 and Imputing Strategy = Global Imputing; Normalization Strategy = Local normalization; Row Selection = Automatic. Relative quantification was performed in Spectronaut for each pairwise comparison using the replicate samples from each condition. The candidates and protein report tables (for protein quantities) were exported from Spectronaut and further data analyses and visualization were performed with RStudio using in-house pipelines and scripts.

### IncuCyte growth curve analysis

Cells were plated at a density of 1.56 x 10^4^ cells/cm^2^ in 96-well-plates. 12 h after plating, the medium was changed by growth medium containing FLI-06, BFA, Thapsigargin (Merck, cat.no. 586005-1MG), Tunicamycin (Merck, cat.no. T7765-1MG) or DMSO as mock treatment. Then, live cell imaging with the IncuCyte S3 microscope (Essen BioScience, cat.no. 4647) was started (t = 0 h) and continued for several days with an interval of 3 h. For each cell line and treatment condition three biological replicates (three wells) were performed and for every experiment appropriate controls (cell lines expressing the empty vector and mock treatment) were included. Per well, four images at distinct positions were captured with the 10x objective. The area occupied by cells (confluence) was determined by the IncuCyte S3 software. Then, the confluence of the four images per well was averaged and delivered as output value. Statistical analysis and visualization were performed with RStudio using in-house scripts.

### Immunofluorescence staining

Cells were cultivated in 8-well µ-Slides (Ibidi, cat.no. 80827) or on glass coverslips. Fixation was performed at RT for 10 min with 4 % formaldehyde (Thermo Fisher Scientific, cat.no. 28906), which was diluted in PBS. After washing 3x with PBS for 5 min, the cells were permeabilized with 0.2 % Triton X-100 in PBS at RT for 5 min. Then, the cells were washed again 3x with PBS. Unspecific binding sites were blocked by incubation with 3 % goat serum (Sigma-Aldrich, cat.no. G9023-10ML) in PBS at RT for 1 h. The cells were probed overnight at 4 °C with the primary antibodies, which were diluted in blocking buffer. After washing 3x with PBS, the cells were incubated at RT for 1 h with fluorescently labelled secondary antibodies and Hoechst-33342 (Sigma-Aldrich, cat.no. B2261-25MG), which were diluted in blocking buffer as well.

### Golgi fragment quantification

Cells were treated with 10 µM of FLI-06 or DMSO (mock treatment, 1:1000) for 24 h before performing immunofluorescence staining for the Golgi protein GolginB1. Microscopic images were acquired with an epifluorescence microscope (Carl Zeiss Microscopy, Axio Imager Z2) and analyzed using Fiji with in-house macros (Behrendt et al., 2021). The number of nuclei (Hoechst-33342) and Golgi fragments per image were quantified and used to calculate the number of Golgi fragments per cell. For each experimental condition 10 microscopic images were quantified per biological replicate. Statistical analysis and visualization were performed with RStudio using in-house scripts.

### VSVG-EYFP transport assay

HeLa cells were co-transfected with pcDNA3.1_VSVG-EYFP_ts045 (Yonemura et al., 2016) and the indicated plasmid / siRNA using Lipofectamine™ 3000 (Thermo Fisher Scientific, cat.no. L3000015) as described by the manufacturer. The cells were then incubated at 40 °C for 14 h. After a 2 h pre-incubation with 10 µM FLI-06 at 40°C, the cells were shifted to 32 °C. Deglycosylation of VSVG-EYFP was performed basically as described (Fassler et al., 2011) but using direct cell lysates instead of immunoprecipitates.

For visualization of VSVG-EYFP-transport w/o ATF6 inhibition, cells were preincubated for the last 30min of the 14h 40° incubation period with 40µg/ml cycloheximide and, where indicated, 10 µM FLI-06, and/or 10 µM CeapinA7 before shifting to 32°C. Before fixation at indicated time points cells were incubated for 20min on ice with anti-VSVG antibody EB0010 to label cell-surface VSVG-EYFP. After fixation in PFA, cells were processed for immunofluorescence using Alexa555 secondary antibody. The surface-to-total VSVG-EYFP ratio was determined using CellProfiler image analysis software. First, nuclei were deteced based on Hoechst 33258 staining by the “IdentifyPrimaryObjects” module. Then the green (for total VSVG-EYFP) and red (for surface-VSVG-EYFP) fluorescence signals were detected using two consecutive “IdentifySecondaryObjects” modules. Afterwards, a filtering step based on a mean fluorescence intensity (MFI) arbitrary threshold was performed, applying the “FilterObjects” module to retain only cells that are positive in both channels. Once the intensities are measured (“MeasureObjectIntensity”), the surface-to-total VSVG ratio was calculated for each single cell by dividing the mean intensity of Alexa555 by the mean intensity of EGFP-VSVG.

### SEAP assay

HeLa cells stably expressing SEAP (Yonemura et al., 2016) were transfected with EGFP-tagged ATL3 (as control) or EGFP-tagged ATF6 and after 24h SEAP assay was performed as described (Yonemura et al., 2016).

### RNA-seq

Total RNA was extracted using RNeasy columns (Qiagen). Sequencing of RNA samples was performed using Illumina’s next-generation sequencing methodology (Bentley et al., 2008). In detail, total RNA was quantified and quality checked using Tapestation 4200 instrument in combination with RNA ScreenTape (both Agilent Technologies). Libraries were prepared from 200 ng of input material (total RNA) using NEBNext Ultra II Directional RNA Library Preparation Kit in combination with NEBNext Poly(A) mRNA Magnetic Isolation Module and NEBNext Multiplex Oligos for Illumina (Unique Dual Index UMI Adaptors RNA) following the manufacturer’s instructions (New England Biolabs). Quantification and quality checked of libraries was done using an Agilent 2100 Bioanalyzer instrument and a DNA 7500 kit (Agilent Technologies). Libraries were pooled and sequenced on a NovaSeq6000 S1 100 cycle run (v1.5 SBS reagents). System run in 101 cycle/single-end/standard loading workflow mode. Sequence information was converted to FASTQ format using bcl2fastq v2.20.0.422.

### CUT&RUN

CUT and RUN experiments were performed as described previously with minor modifications (Kim et al., 2022). Briefly, for each CUT and RUN reaction 200,000 cells were trypsinized, washed, resuspended in 100 µl wash buffer (20 mM HEPES, pH7.5, 150 mM NaCl, 0.5 mM Spermidine) and bound to 10 µl activated Concanavalin A magnetic beads (Polysciences) for 10 min at room temperature. The cells were then incubated with antibodies (ATF6: Abcam #227830; XBP1: Cell Signaling #40435) diluted in 100 µl antibody buffer (wash buffer + 0.01% digitonine and 2 mM EDTA) overnight at 4°C. Rabbit IgG was used as negative control. After antibody incubation, the beads were washed in digitonin wash buffer (wash buffer + 0.01% digitonin) and incubated 1 h at 4°C with 1 µg/ml protein A/G Micrococcal Nuclease fusion protein (pA/G MNase). After washing with digitonin wash buffer, beads were rinsed with low salt buffer (20 mM HEPES, pH7.5, 0.5 mM Spermidine, 0.01% digitonine), resuspended in 200 µl digitonin wash buffer and placed at 4°C. To initiate cleavage, 4 µl of 0.1M CaCl2 was added to the beads. After 1h, the buffer was removed and reactions were stopped by adding 200 µl STOP buffer (170 mM NaCl, 20 mM EGTA, 0.01% digitonin, 50 µg/ml RNAse A) and the samples were incubated 30 min at 37°C to digest the RNA and release the DNA. The samples were then treated with proteinase K for 1 h at 50°C and the DNA was purified using Phenol/Chloroform/Isoamyl alcohol. After EtOH precipitation, the DNA pellet was resuspended in 0.1 X TE and used for DNA library generation with the NEBNext® Ultra™ II DNA Library Prep Kit for Illumina® (New England Biolabs). Adaptor ligation was performed with 1:25 diluted adaptor and 15 cycles were used for library amplification using dual indices (NEB dual index kit). Paired-end 2×25 bp sequencing was performed on a NextSeq500 Illumina Sequencer.

### CUT&RUN analysis

Cutadapt was used for adapter removal and quality trimming. The carry-over *E. coli* DNA from pAG-MNase purification was used as spike-in control, mapping was performed to hg19 and a human repeat-masked *E. coli* genome by Bowtie2. Paired-end reads mapped to hg19 with inserts <120 bp were extracted using alignment Sieve (deepTools). A scaling factor was inferred by:

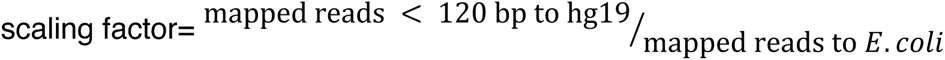

The scaling factor was used to generate spike-in normalized bigWig files by bamCoverage (deepTools). The bigwig files were further normalized to sequencing depth by CPM. For peak calling, Genrich was used with a 1e-9 q-value cutoff. For quantitative analyses, the spike-normalized bigWig files were used in computeMatrix reference-point (DeepTools), e.g. using peaks as reference point. The matrix output of computeMatrix, was then used for further analyses in R.

#### RNA-Sequencing analysis

Adapter removal, size selection (reads > 25 nt) and quality filtering (Phred score > 43) of FASTQ files were performed with Cutadapt. Reads were then aligned to human genome (hg19) using Bowtie2 (v2.2.9) with default settings. Read count extraction was performed in R using countOverlaps (GenomicRanges).

Differential gene expression analysis was done with DESeq2 (v3.26.8) using default parameters. Cluster analysis of YAP target genes was performed by MFuzz in R using three clusters.

### Statistics

All statistical tests were performed using R. The specific statistical test and number of biological replicates are always indicated in the respective figure legend.

## Data availability

Sequencing data were deposited at GEO repository, accession numbers pending.

## Acknowledgements

BvE was supported by grants from the BMBF (16GW0271K), DFG (EY 120/4-1), and the German Cancer Aid (Deutsche Krebshilfe; 70113138). CK was supported by grants from the DFG (KA1751/8-1; KA1751/9-1). The FLI is a member of the Leibniz Association and is financially supported by the Federal Government of Germany and the State of Thuringia. We are grateful to all donors of Addgene plasmids. We thank the core facilities at the FLI CF imaging, CF Sequencing and CF Proteomics.

## Author contributions

CK and BvE conceived the project and designed experiments. PC, YY, LB, AM, TM and JM performed experiments and analyzed the data. MS and CG performed the LOF screen, KS analyzed data, NN contributed research tools. CK and BvE wrote the paper with help of all authors. All authors discussed the results and implications at all stages.

---

**Suppl. Fig. 1: Validation of additional hits of the CRISPR_a_ SAM screen for BFA / FLI-06 resistance.**

**a, b**) Scatter plot (left) and violin plot (right) of sgRNA frequency of the CRISPR_a_ SAM screen for BFA resistance in the two clonal MCF10A-SAM cell lines #9 (**a**) and #28 (**b**). *GBF1*-targeting sgRNAs are highlighted (blue). **c-h**) Validation of *ATF3*, *EPHX1*, *PCNX2* and *PRRC1* as potential BFA targets. qPCR expression analysis (**c-f**) of the depicted genes and IncuCyte growth curve analysis upon BFA treatment (**g, h**) in MCF10A-SAM cells stably expressing the indicated SAM-sgRNAs or non-targeting (NT) sgRNAs. Displayed is the confluency over time. **i, j**) Scatter plot (left) and violin plot (right) of sgRNA frequency of the CRISPR_a_ SAM screen for FLI-06 resistance in the two clonal MCF10A-SAM cell lines #9 (**i**) and #28 (**j**). *GOLT1A*- / *GOLT1B*-targeting sgRNAs are highlighted (blue / red). **k-o**) Validation of *GPR89B*, *RNF13*, *TMEM52* and *PRRC1* as potential FLI-06 targets. qPCR expression analysis (**k-m**) of the depicted genes and IncuCyte growth curve analysis upon FLI-06 treatment (**n, o**) in MCF10A-SAM cells stably expressing the indicated SAM-sgRNAs or non-targeting (NT) sgRNAs. **a, b, i, j**: Each dot represents one sgRNA. Statistical testing: one-sample Wilcox test against the hypothetical mean of zero. **c-h, k-o**: Data are shown as mean of three biological replicates and error bars represent SEM. Statistical testing: one-way-ANOVA test with Tukey HSD post-hoc test.

**Suppl. Fig. 2: Overexpression of *GOLT1A* renders cells FLI-06 resistant, while *GOLT1B* overexpression sensitizes cells to FLI-06.**

**a**) Protein alignment of human GOT1A and GOT1B. Transmembrane domains (TMD) were annotated according to UniProt (release 04/2021). **b, c**) IncuCyte growth curve analysis upon FLI-06 treatment (20 µM, **b**) or DMSO (**c**) in MCF10A-SAM clone #9 cells stably expressing *GOLT1A*- or non-targeting (NT) sgRNAs. **d, e**) qPCR expression analysis (**d**) of *GOLT1B* and IncuCyte growth curve analysis (**e**) upon FLI-06 treatment (5 µM) in MCF10A-SAM clone #9 cells stably expressing *GOLT1B*- or non-targeting sgRNAs. For qPCR, *B2M* was used as housekeeping gene. **f**) MCF10A-SAM clone #9 cells stably expressing *GOLT1A*- *GOLT1B*- or non-targeting sgRNAs were treated for 24 h with 10 µM FLI-06 or DMSO as control, fixed and processed for immunofluorescence staining against the Golgi marker GolginB1 (green). Counterstaining with Hoechst-33342 (magenta) visualizes nuclei. Representative images of three biological replicates are shown. Scale bar: 10 µm. Inserts (white square, scale bar: 5 µm) show higher magnification of the respective area. **g**) Quantification of Golgi-fragments based on immunofluorescence images of **f**). **h,i**) Lentiviral expression in MCF10A cells of N- or C-terminally tagged *GOLT1A* and *GOLT1B* constructs, respectively. Immunoblotting analysis of stably expressed *GOLT1A* (**h**) and *GOLT1B* (**i**). Vinculin served as loading control. **j**, **k**) IncuCyte growth curve analysis upon FLI-06 treatment in MCF10A cells stably expressing the indicated *GOLT1A*/*B* constructs. Data are shown as mean of three biological replicates and error bars represent SEM. Statistical testing: one-way-ANOVA test with Tukey HSD post-hoc test.

**Suppl. Fig. 3: GOT1A and GOT1B localize to the early secretory pathway.**

**a-b**) Immunofluorescence staining of MCF10A cells stably expressing *Flag-GOLT1A*(**a**) or *GOLT1B-Flag* (**b**). Cells were stained with α-Flag-M2 (red) to label GOT1A or GOT1B. Additionally, compartments of the secretory pathway were labeled (green) by co-staining with α-RTN4 (ER), α-SEC16A (ERES), α-ERGIC53 (ERGIC), α-GolginB1 (Golgi), α-βCOP (COPI vesicle) and α-YIPF5. Counterstaining with Hoechst-33342 (blue) visualizes nuclei. Arrowheads indicate areas of co-localization of GOT1A or GOT1B with the respective compartments of the secretory pathway or YIPF5, respectively. Representative images of three biological replicates are shown. Scale bar: 10 µm. Inserts (white square, scale bar: 5 µm) show higher magnification of the respective area. **c, d**) IncuCyte growth curve analysis upon Tunicamycin (**c**) or Thapsigargin (**d**) treatment in stable MCF10A cells expressing *Flag-GOLT1A* and *GOLT1B-Flag*. Data are shown as mean of three biological replicates and error bars represent SEM. Statistical testing: one-way-ANOVA test with Tukey HSD post-hoc test.

**Suppl. Fig. 4: Protein-protein interactome of GOT1A and GOT1B.**

**a-d**) Volcano plot of IP-MS analysis of MCF10A cells stably expressing *Flag-GOLT1A* (**a, b**) or *GOLT1B-Flag* (**c, d**). The cells were pretreated for 14 h with DMSO (**a, c**) or 10 µM FLI-06 (**b, d**) and IP was performed using Flag-antibodies. The Log_2_FC was calculated in comparison with empty vector expressing cells. Significant protein-protein interactors were defined as Log_2_FC ≥ 0.58 and q-value ≤ 0.05 (marked with dashed lines). n = four biological replicates. The bait protein (Flag-GOT1A / GOT1B-Flag) and validated interactors are labeled. **e, f**) Venn diagram showing the overlap of GOT1A-and GOT1B interactors upon DMSO (**e**) or FLI-06 (**f**) treatment. **g-j**) qPCR expression analysis of *HERPUD1* (**g,i**) and *P4HB* (**h,j**) in MCF10A cells stably expressing *Flag-GOLT1A* (**g,h**) or siGOLT1B (**i,j**) transfected. Cells were treated with Thapsigargin (100 nM) for 14 h. Data are shown as mean of three biological replicates and error bars represent SEM. Statistical testing: one-way-ANOVA test with Tukey HSD post-hoc test.

**Suppl. Figure 5:** Control for cycloheximide and overexpression of ATF6 does not block ER export.

**a)** Cells were treated for 2h with 10 µM cycloheximide or DMSO, fixed and processed for immunofluorescence with antibodies against the short-lived protein c-MYC. Scalebar 10μm. **b)** Hela cells stably expressing SEAP were transfected with ATL3-GFP as an unrelated membrane protein or GFP-ATF6, lysed after 24h, processed for Western Blot and probed with anti-ATF6 antibodies. **c)** Quantification of SEAP secretion. Media of cells from b) were collected and a SEAP activity assay performed. Data are normalized to ATL3-GFP cells as control. Two-way one-sample Welch T-test. **d)** Hela cells were transfected with GFP-ATF6 and VSVG-mCherry and incubated overnight at 40°C. After a 2h chase at 32°C, cells were put on ice, the surface VSVG-mCherry stained with an antibody against VSVG before fixation and labeling with secondary Cy5-labelled antibody and the cells imaged by fluorescence microscopy. Scalebar 10μm.

## References

Adolf F, Rhiel M, Hessling B, Gao Q, Hellwig A, Bethune J, Wieland FT (2019) Proteomic Profiling of Mammalian COPII and COPI Vesicles. Cell Rep 26: 250–265 e5

Aridor M (2018) COPII gets in shape: Lessons derived from morphological aspects of early secretion. Traffic 19: 823–839

Barlowe C, Helenius A (2016) Cargo Capture and Bulk Flow in the Early Secretory Pathway. Annu Rev Cell Dev Biol 32: 197–222

Behrendt L, Hoischen C, Kaether C (2021) Disease-causing mutated ATLASTIN 3 is excluded from distal axons and reduces axonal autophagy. Neurobiol Dis 155: 105400

Bentley DR, Balasubramanian S, Swerdlow HP, Smith GP, Milton J, Brown CG, Hall KP, Evers DJ, Barnes CL, Bignell HR, Boutell JM, Bryant J, Carter RJ, Keira Cheetham R, Cox AJ, Ellis DJ, Flatbush MR, Gormley NA, Humphray SJ, Irving LJ et al. (2008) Accurate whole human genome sequencing using reversible terminator chemistry. Nature 456: 53–9

Centonze FG, Reiterer V, Nalbach K, Saito K, Pawlowski K, Behrends C, Farhan H (2019) LTK is an ER-resident receptor tyrosine kinase that regulates secretion. Journal of Cell Biology 218: 2470–2480

Claude A, Zhao BP, Kuziemsky CE, Dahan S, Berger SJ, Yan JP, Armold AD, Sullivan EM, Melancon P (1999) GBF1: A novel Golgi-associated BFA-resistant guanine nucleotide exchange factor that displays specificity for ADP-ribosylation factor 5. J Cell Biol 146: 71–84

Conchon S, Cao X, Barlowe C, Pelham HR (1999) Got1p and Sft2p: membrane proteins involved in traffic to the Golgi complex. EMBO J 18: 3934–46

Ellgaard L, Helenius A (2003) Quality control in the endoplasmic reticulum. Nat Rev Mol Cell Biol 4: 181–91

Elster D, Tollot M, Schlegelmilch K, Ori A, Rosenwald A, Sahai E, von Eyss B (2018) TRPS1 shapes YAP/TEAD-dependent transcription in breast cancer cells. Nat Commun 9: 3115

Fassler M, Li X, Kaether C (2011) Polar transmembrane-based amino acids in presenilin 1 are involved in endoplasmic reticulum localization, Pen2 protein binding, and gamma-secretase complex stabilization. The Journal of biological chemistry 286: 38390–6

Foufelle F, Fromenty B (2016) Role of endoplasmic reticulum stress in drug-induced toxicity. Pharmacol Res Perspect 4: e00211

Fujiwara T, Oda K, Yokota S, Takatsuki A, Ikehara Y (1988) Brefeldin A causes disassembly of the Golgi complex and accumulation of secretory proteins in the endoplasmic reticulum. J Biol Chem 263: 18545–52

Gallagher CM, Walter P (2016) Ceapins inhibit ATF6alpha signaling by selectively preventing transport of ATF6alpha to the Golgi apparatus during ER stress. eLife 5

Gallione CJ, Rose JK (1985) A single amino acid substitution in a hydrophobic domain causes temperature-sensitive cell-surface transport of a mutant viral glycoprotein. Journal of virology 54: 374–82

Gomez-Navarro N, Miller E (2016) Protein sorting at the ER-Golgi interface. J Cell Biol 215: 769–778

Gunaratne GS, Rebbeck RT, McGurran LM, Yan Y, Arzua T, Frolkis T, Sprague DJ, Bai X, Cornea RL, Walseth TF, Marchant JS (2022) Identification of a dihydropyridine scaffold that blocks ryanodine receptors. iScience 25: 103706

Haze K, Yoshida H, Yanagi H, Yura T, Mori K (1999) Mammalian transcription factor ATF6 is synthesized as a transmembrane protein and activated by proteolysis in response to endoplasmic reticulum stress. Mol Biol Cell 10: 3787–99

Heidtman M, Chen CZ, Collins RN, Barlowe C (2003) A role for Yip1p in COPII vesicle biogenesis. J Cell Biol 163: 57–69

Hein MY, Hubner NC, Poser I, Cox J, Nagaraj N, Toyoda Y, Gak IA, Weisswange I, Mansfeld J, Buchholz F, Hyman AA, Mann M (2015) A human interactome in three quantitative dimensions organized by stoichiometries and abundances. Cell 163: 712–23

Hetz C, Zhang KZ, Kaufman RJ (2020) Mechanisms, regulation and functions of the unfolded protein response. Nat Rev Mol Cell Bio 21: 421–438

Higashio H, Kohno K (2002) A genetic link between the unfolded protein response and vesicle formation from the endoplasmic reticulum. Biochem Bioph Res Co 296: 568–574

Ikeda K, Horie-Inoue K, Ueno T, Suzuki T, Sato W, Shigekawa T, Osaki A, Saeki T, Berezikov E, Mano H, Inoue S (2015) miR-378a-3p modulates tamoxifen sensitivity in breast cancer MCF-7 cells through targeting GOLT1A. Scientific reports 5: 13170

Jost M, Weissman JS (2018) CRISPR Approaches to Small Molecule Target Identification. ACS Chem Biol 13: 366–375

Joung J, Konermann S, Gootenberg JS, Abudayyeh OO, Platt RJ, Brigham MD, Sanjana NE, Zhang F (2017) Genome-scale CRISPR-Cas9 knockout and transcriptional activation screening. Nature protocols 12: 828–863

Kim KM, Mura-Meszaros A, Tollot M, Krishnan MS, Grundl M, Neubert L, Groth M, Rodriguez-Fraticelli A, Svendsen AF, Campaner S, Andreas N, Kamradt T, Hoffmann S, Camargo FD, Heidel FH, Bystrykh LV, de Haan G, von Eyss B (2022) Taz protects hematopoietic stem cells from an aging-dependent decrease in PU.1 activity. Nat Commun 13: 5187

Klausner RD, Donaldson JG, Lippincott-Schwartz J (1992) Brefeldin A: insights into the control of membrane traffic and organelle structure. J Cell Biol 116: 1071–80

Konermann S, Brigham MD, Trevino AE, Joung J, Abudayyeh OO, Barcena C, Hsu PD, Habib N, Gootenberg JS, Nishimasu H, Nureki O, Zhang F (2015) Genome-scale transcriptional activation by an engineered CRISPR-Cas9 complex. Nature 517: 583–8

Konig R, Chiang CY, Tu BP, Yan SF, DeJesus PD, Romero A, Bergauer T, Orth A, Krueger U, Zhou Y, Chanda SK (2007) A probability-based approach for the analysis of large-scale RNAi screens. Nat Methods 4: 847–9

Krämer A, Mentrup T, Kleizen B, Rivera-Milla E, Reichenbach D, Enzensperger C, Nohl R, Tauscher E, Gorls H, Ploubidou A, Englert C, Werz O, Arndt HD, Kaether C (2013) Small molecules intercept Notch signaling and the early secretory pathway. Nat Chem Biol 9: 731–8

Kranjc T, Dempsey E, Cagney G, Nakamura N, Shields DC, Simpson JC (2017) Functional characterisation of the YIPF protein family in mammalian cells. Histochem Cell Biol 147: 439–451

Lorente-Rodriguez A, Heidtman M, Barlowe C (2009) Multicopy suppressor analysis of thermosensitive YIP1 alleles implicates GOT1 in transport from the ER. J Cell Sci 122: 1540–50

Ma W, Goldberg E, Goldberg J (2017) ER retention is imposed by COPII protein sorting and attenuated by 4-phenylbutyrate. eLife 6

Malis Y, Hirschberg K, Kaether C (2022) Hanging the coat on a collar: Same function but different localization and mechanism for COPII. Bioessays 44: e2200064

McCaughey J, Stephens DJ (2018) COPII-dependent ER export in animal cells: adaptation and control for diverse cargo. Histochem Cell Biol 150: 119–131

Mezzacasa A, Helenius A (2002) The transitional ER defines a boundary for quality control in the secretion of tsO45 VSV glycoprotein. Traffic 3: 833–49

Misumi Y, Misumi Y, Miki K, Takatsuki A, Tamura G, Ikehara Y (1986) Novel blockade by brefeldin A of intracellular transport of secretory proteins in cultured rat hepatocytes. J Biol Chem 261: 11398–403

Raote I, Saxena S, Malhotra V (2023) Sorting and Export of Proteins at the Endoplasmic Reticulum. Cold Spring Harbor perspectives in biology 15

Ron D, Walter P (2007) Signal integration in the endoplasmic reticulum unfolded protein response. Nat Rev Mol Cell Bio 8: 519–529

Saenz JB, Sun WJ, Chang JW, Li J, Bursulaya B, Gray NS, Haslam DB (2009) Golgicide A reveals essential roles for GBF1 in Golgi assembly and function. Nat Chem Biol 5: 157–65

Sato M, Sato K, Nakano A (2002) Evidence for the intimate relationship between vesicle budding from the ER and the unfolded protein response. Biochem Bioph Res Co 296: 560–567

Schweizer A, Rohrer J, Hauri HP, Kornfeld S (1994) Retention of p63 in an ER-Golgi intermediate compartment depends on the presence of all three of its domains and on its ability to form oligomers. J Cell Biol 126: 25–39

Shaffer AL, Shapiro-Shelef M, Iwakoshi NN, Lee AH, Qian SB, Zhao H, Yu X, Yang LM, Tan BK, Rosenwald A, Hurt EM, Petroulakis E, Sonenberg N, Yewdell JW, Calame K, Glimcher LH, Staudt LM (2004) XBP1, downstream of Blimp-1, expands the secretory apparatus and other organelles, and increases protein synthesis in plasma cell differentiation. Immunity 21: 81–93

Shaheen A (2018) Effect of the unfolded protein response on ER protein export: a potential new mechanism to relieve ER stress. Cell Stress Chaperones 23: 797–806

Shalem O, Sanjana NE, Hartenian E, Shi X, Scott DA, Mikkelsen TS, Heckl D, Ebert BL, Root DE, Doench JG, Zhang F (2014) Genome-scale CRISPR-Cas9 knockout screening in human cells. Science 343: 84–7

Shomron O, Nevo-Yassaf I, Aviad T, Yaffe Y, Zahavi EE, Dukhovny A, Perlson E, Brodsky I, Yeheskel A, Pasmanik-Chor M, Mironov A, Beznoussenko GV, Mironov AA, Sklan EH, Patterson GH, Yonemura Y, Sannai M, Kaether C, Hirschberg K (2021) COPII collar defines the boundary between ER and ER exit site and does not coat cargo containers. J Cell Biol 220

Sriburi R, Bommiasamy H, Buldak GL, Robbins GR, Frank M, Jackowski S, Brewer JW (2007) Coordinate regulation of phospholipid biosynthesis and secretory pathway gene expression in XBP-1 (S)-induced endoplasmic reticulum biogenesis. Journal of Biological Chemistry 282: 7024–7034

Subramanian A, Capalbo A, Iyengar NR, Rizzo R, di Campli A, Di Martino R, Lo Monte M, Beccari AR, Yerudkar A, del Vecchio C, Glielmo L, Turacchio G, Pirozzi M, Kim SG, Henklein P, Cancino J, Parashuraman S, Diviani D, Fanelli F, Sallese M et al. (2019) Auto-regulation of Secretory Flux by Sensing and Responding to the Folded Cargo Protein Load in the Endoplasmic Reticulum. Cell 176: 1461-+

Tang BL, Ong YS, Huang B, Wei S, Wong ET, Qi R, Horstmann H, Hong W (2001) A membrane protein enriched in endoplasmic reticulum exit sites interacts with COPII. J Biol Chem 276: 40008–17

Wang Y, Liu F, Ren Y, Wang Y, Liu X, Long W, Wang D, Zhu J, Zhu X, Jing R, Wu M, Hao Y, Jiang L, Wang C, Wang H, Bao Y, Wan J (2016) GOLGI TRANSPORT 1B Regulates Protein Export from the Endoplasmic Reticulum in Rice Endosperm Cells. Plant Cell 28: 2850–2865

Weigel AV, Chang CL, Shtengel G, Xu CS, Hoffman DP, Freeman M, Iyer N, Aaron J, Khuon S, Bogovic J, Qiu W, Hess HF, Lippincott-Schwartz J (2021) ER-to-Golgi protein delivery through an interwoven, tubular network extending from ER. Cell 184: 2412–2429 e16

Yamaji R, Adamik R, Takeda K, Togawa A, Pacheco-Rodriguez G, Ferrans VJ, Moss J, Vaughan M (2000) Identification and localization of two brefeldin A-inhibited guanine nucleotide-exchange proteins for ADP-ribosylation factors in a macromolecular complex. Proc Natl Acad Sci U S A 97: 2567–72

Ye J, Rawson RB, Komuro R, Chen X, Davé UP, Prywes R, Brown MS, Goldstein JL (2000) ER stress induces cleavage of membrane-bound ATF6 by the same proteases that process SREBPs. Mol Cell 6: 1355–64

Yonemura Y, Li X, Muller K, Kramer A, Atigbire P, Mentrup T, Feuerhake T, Kroll T, Shomron O, Nohl R, Arndt HD, Hoischen C, Hemmerich P, Hirschberg K, Kaether C (2016) Inhibition of cargo export at ER exit sites and the trans-Golgi network by the secretion inhibitor FLI-06. J Cell Sci 129: 3868–3877

Yoshida Y, Suzuki K, Yamamoto A, Sakai N, Bando M, Tanimoto K, Yamaguchi Y, Sakaguchi T, Akhter H, Fujii G, Yoshimura S, Ogata S, Sohda M, Misumi Y, Nakamura N (2008) YIPF5 and YIF1A recycle between the ER and the Golgi apparatus and are involved in the maintenance of the Golgi structure. Exp Cell Res 314: 3427–43

Zanetti G, Pahuja KB, Studer S, Shim S, Schekman R (2012) COPII and the regulation of protein sorting in mammals. Nature cell biology 14: 20–8

Zhang L, Hu R, Cheng Y, Wu X, Xi S, Sun Y, Jiang H (2017) Lidocaine inhibits the proliferation of lung cancer by regulating the expression of GOLT1A. Cell Prolif 50

