## Supplemental figures and info for "A YIPF5-GOT1A/B complex directs a transcription-independent function of ATF6 in ER export"

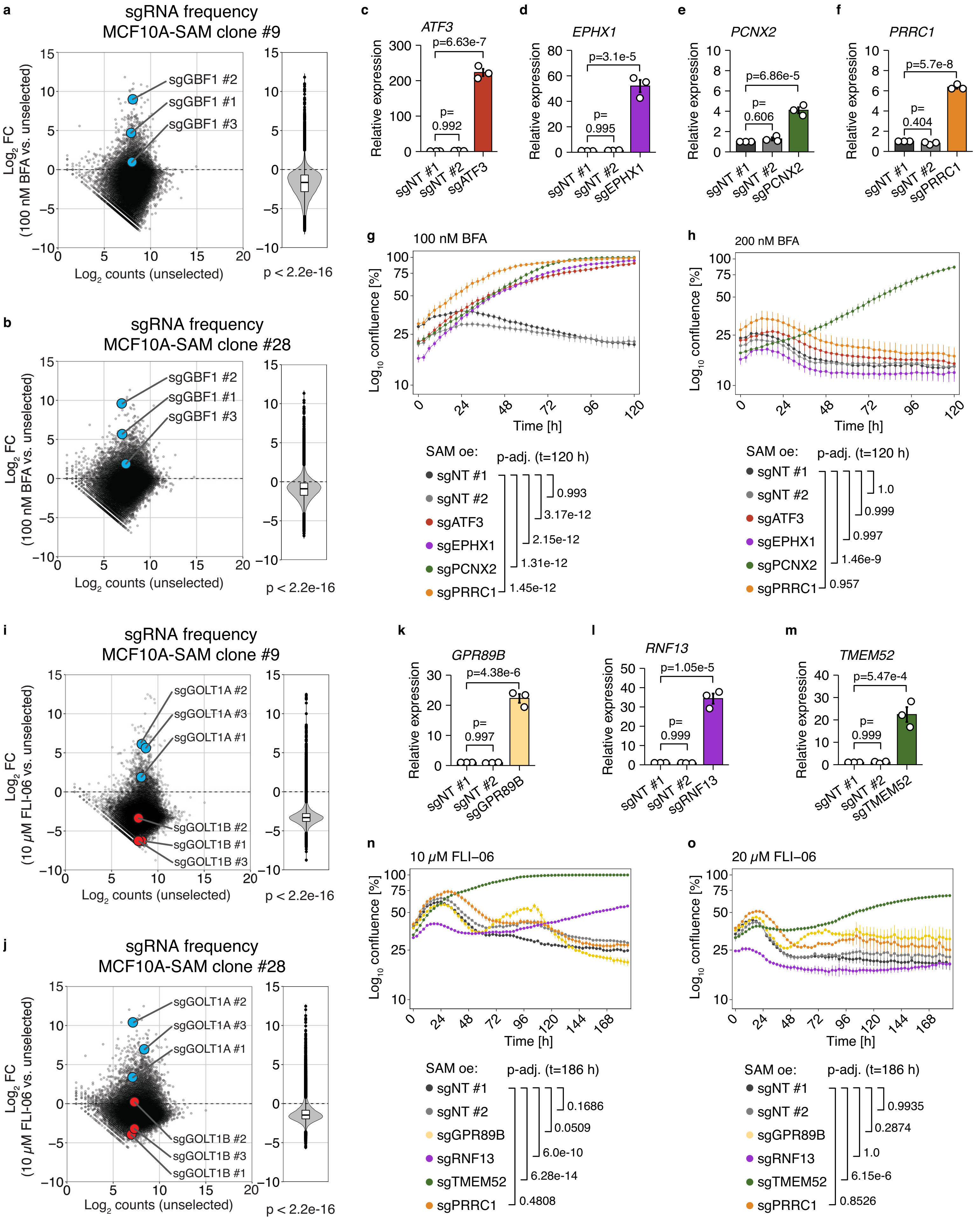

a
sgRNA frequency
MCF10A-SAM clone #9
sgGBF1 #2
sgGBF1 #1
sgGBF1 #3
Log2 FC (100 nM BFA vs. unselected)
Log2 counts (unselected)
p < 2.2e-16
b
sgRNA frequency
MCF10A-SAM clone #28
sgGBF1 #2
sgGBF1 #1
sgGBF1 #3
Log2 FC (100 nM BFA vs. unselected)
Log2 counts (unselected)
p < 2.2e-16
c
ATF3
p=6.63e-7
p= 0.992
Relative expression
sgNT #1
sgNT #2
sgATF3
d
EPHX1
p=3.1e-5
p= 0.995
Relative expression
sgNT #1
sgNT #2
sgEPHX1
e
PCNX2
p=6.86e-5
p= 0.606
Relative expression
sgNT #1
sgNT #2
sgPCNX2
f
PRRC1
p=5.7e-8
p= 0.404
Relative expression
sgNT #1
sgNT #2
sgPRRC1
g
100 nM BFA
Log10 confluence [%]
Time [h]
SAM oe:
p-adj. (t=120 h)
sgNT #1
sgNT #2
sgATF3
sgEPHX1
sgPCNX2
sgPRRC1
0.993
3.17e-12
2.15e-12
1.31e-12
1.45e-12
h
200 nM BFA
Log10 confluence [%]
Time [h]
SAM oe:
p-adj. (t=120 h)
sgNT #1
sgNT #2
sgATF3
sgEPHX1
sgPCNX2
sgPRRC1
1.0
0.999
0.997
1.46e-9
0.957
i
sgRNA frequency
MCF10A-SAM clone #9
sgGOLT1A #2
sgGOLT1A #3
sgGOLT1A #1
sgGOLT1B #2
sgGOLT1B #1
sgGOLT1B #3
Log2 FC (10 μM FLI-06 vs. unselected)
Log2 counts (unselected)
p < 2.2e-16
j
sgRNA frequency
MCF10A-SAM clone #28
sgGOLT1A #2
sgGOLT1A #3
sgGOLT1A #1
sgGOLT1B #2
sgGOLT1B #3
sgGOLT1B #1
Log2 FC (10 μM FLI-06 vs. unselected)
Log2 counts (unselected)
p < 2.2e-16
k
GPR89B
p=4.38e-6
p= 0.997
Relative expression
sgNT #1
sgNT #2
sgGPR89B
l
RNF13
p=1.05e-5
p= 0.999
Relative expression
sgNT #1
sgNT #2
sgRNF13
m
TMEM52
p=5.47e-4
p= 0.999
Relative expression
sgNT #1
sgNT #2
sgTMEM52
n
10 μM FLI-06
Log10 confluence [%]
Time [h]
SAM oe:
p-adj. (t=186 h)
sgNT #1
sgNT #2
sgGPR89B
sgRNF13
sgTMEM52
sgPRRC1
0.1686
0.0509
6.0e-10
6.28e-14
0.4808
o
20 μM FLI-06
Log10 confluence [%]
Time [h]
SAM oe:
p-adj. (t=186 h)
sgNT #1
sgNT #2
sgGPR89B
sgRNF13
sgTMEM52
sgPRRC1
0.9935
0.2874
1.0
6.15e-6
0.8526

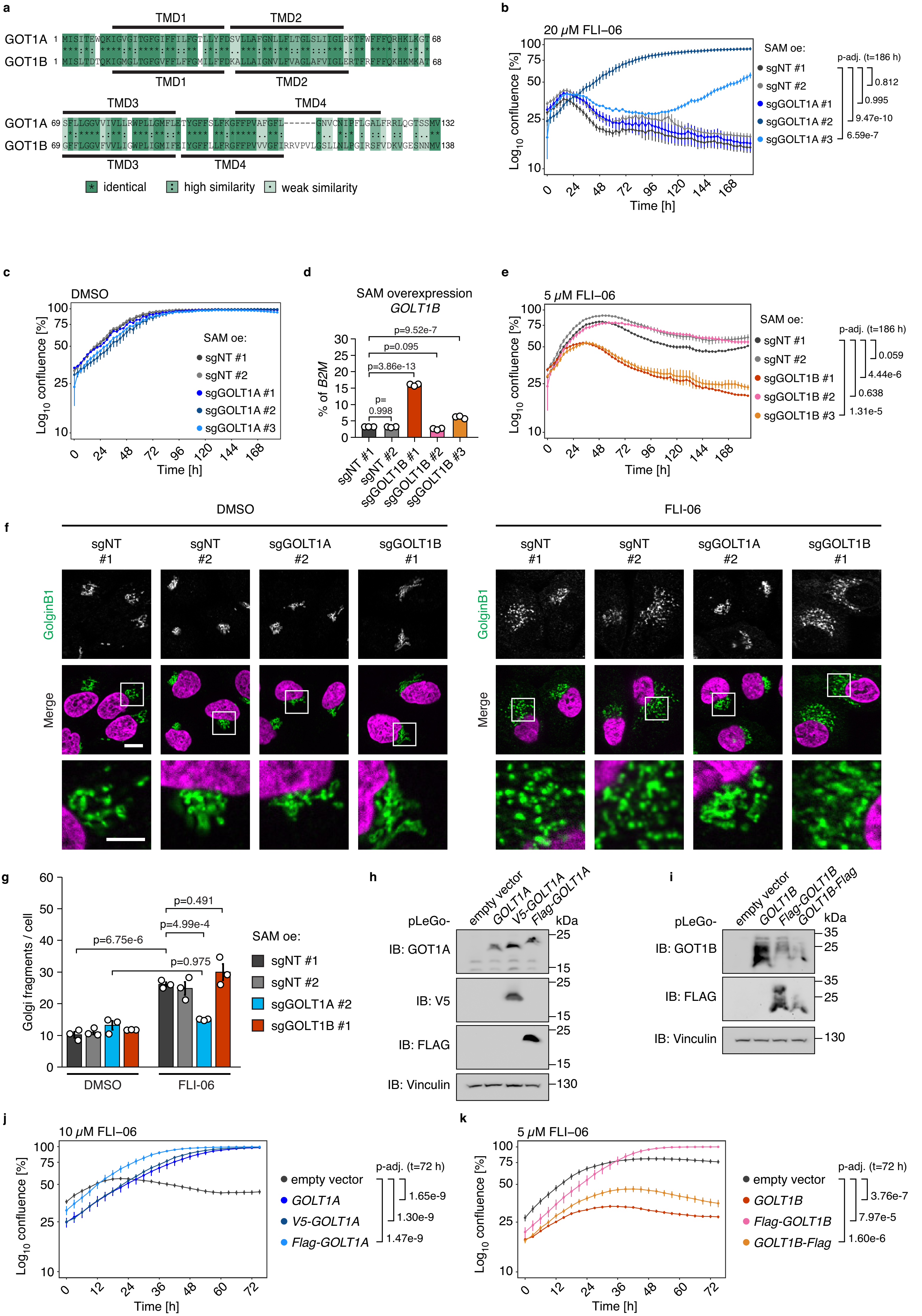

a
TMD1
TMD2
GOT1A 1
GOT1B 1
68
TMD3
TMD4
GOT1A 69
GOT1B 69
132
138
identical
high similarity
weak similarity
b
20 μM FLI–06
Log10 confluence [%]
Time [h]
SAM oe:
sgNT #1
sgNT #2
sgGOLT1A #1
sgGOLT1A #2
sgGOLT1A #3
p-adj. (t=186 h)
0.812
0.995
9.47e-10
6.59e-7
c
DMSO
d
SAM overexpression
GOLT1B
% of B2M
p=9.52e-7
p=0.095
p=3.86e-13
p=
0.998
sgNT #1
sgNT #2
sgGOLT1B #1
sgGOLT1B #2
sgGOLT1B #3
e
5 μM FLI–06
0.059
4.44e-6
0.638
1.31e-5
DMSO
FLI-06
f
GolginB1
Merge
g
Golgi fragments / cell
p=6.75e-6
p=4.99e-4
p=0.491
p=0.975
h
pLeGo-
empty vector
GOLT1A
V5-GOLT1A
Flag-GOLT1A
kDa
IB: GOT1A
IB: V5
IB: FLAG
IB: Vinculin
i
GOLT1B
Flag-GOLT1B
GOLT1B-Flag
IB: GOT1B
j
10 μM FLI–06
p-adj. (t=72 h)
1.65e-9
1.30e-9
1.47e-9
k
3.76e-7
7.97e-5
1.60e-6

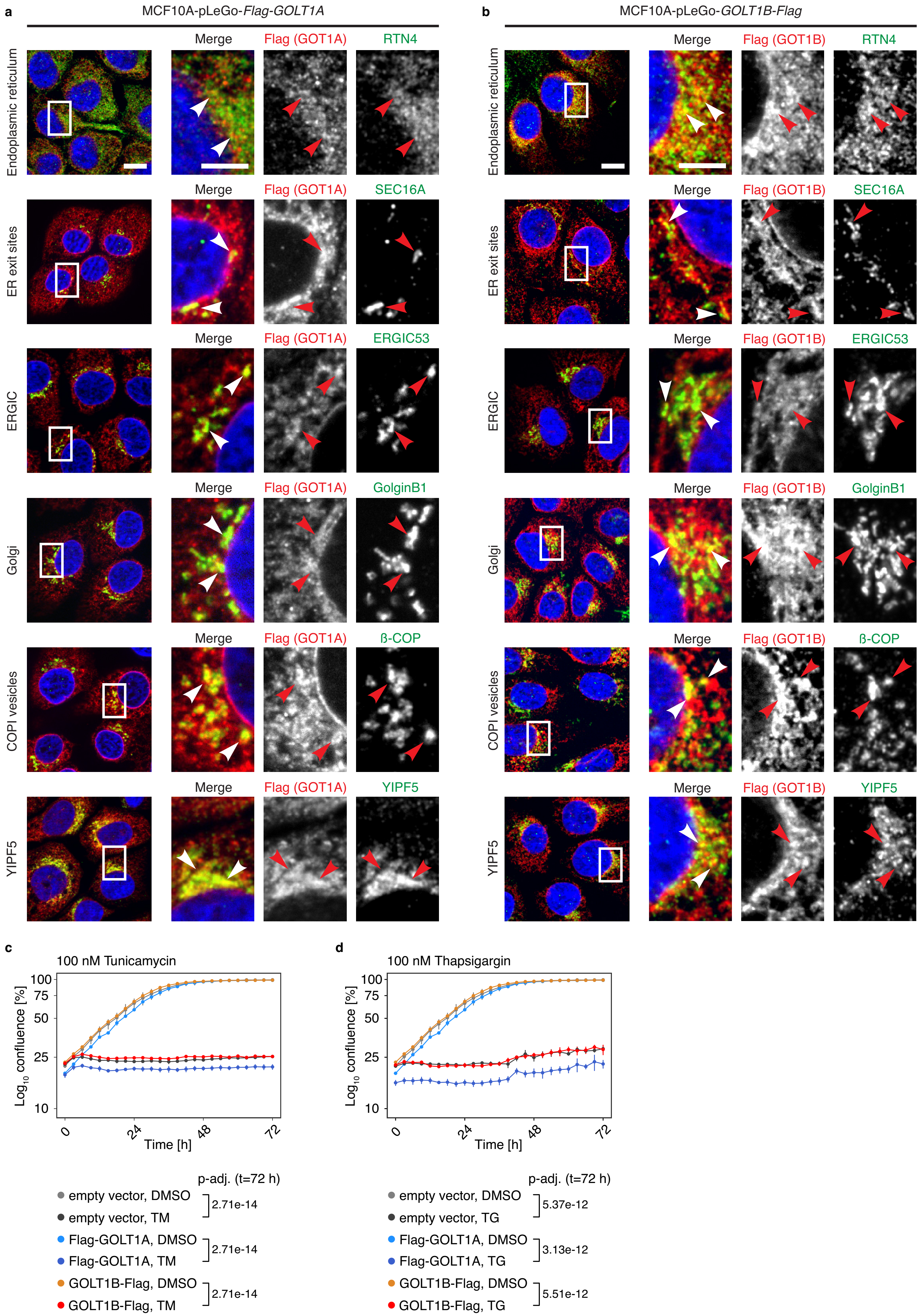
a
MCF10A-pLeGo-Flag-GOLT1A
b
MCF10A-pLeGo-GOLT1B-Flag
Endoplasmic reticulum
Merge
Flag (GOT1A)
RTN4
Flag (GOT1B)
ER exit sites
SEC16A
ERGIC
ERGIC53
Golgi
GolginB1
COPI vesicles
ß-COP
YIPF5
c
100 nM Tunicamycin
d
100 nM Thapsigargin
Log10 confluence [%]
100
75
50
25
10
0
24
48
72
Time [h]
p-adj. (t=72 h)
empty vector, DMSO
empty vector, TM
Flag-GOLT1A, DMSO
Flag-GOLT1A, TM
GOLT1B-Flag, DMSO
GOLT1B-Flag, TM
2.71e-14
2.71e-14
2.71e-14
empty vector, DMSO
empty vector, TG
Flag-GOLT1A, DMSO
Flag-GOLT1A, TG
GOLT1B-Flag, DMSO
GOLT1B-Flag, TG
5.37e-12
3.13e-12
5.51e-12

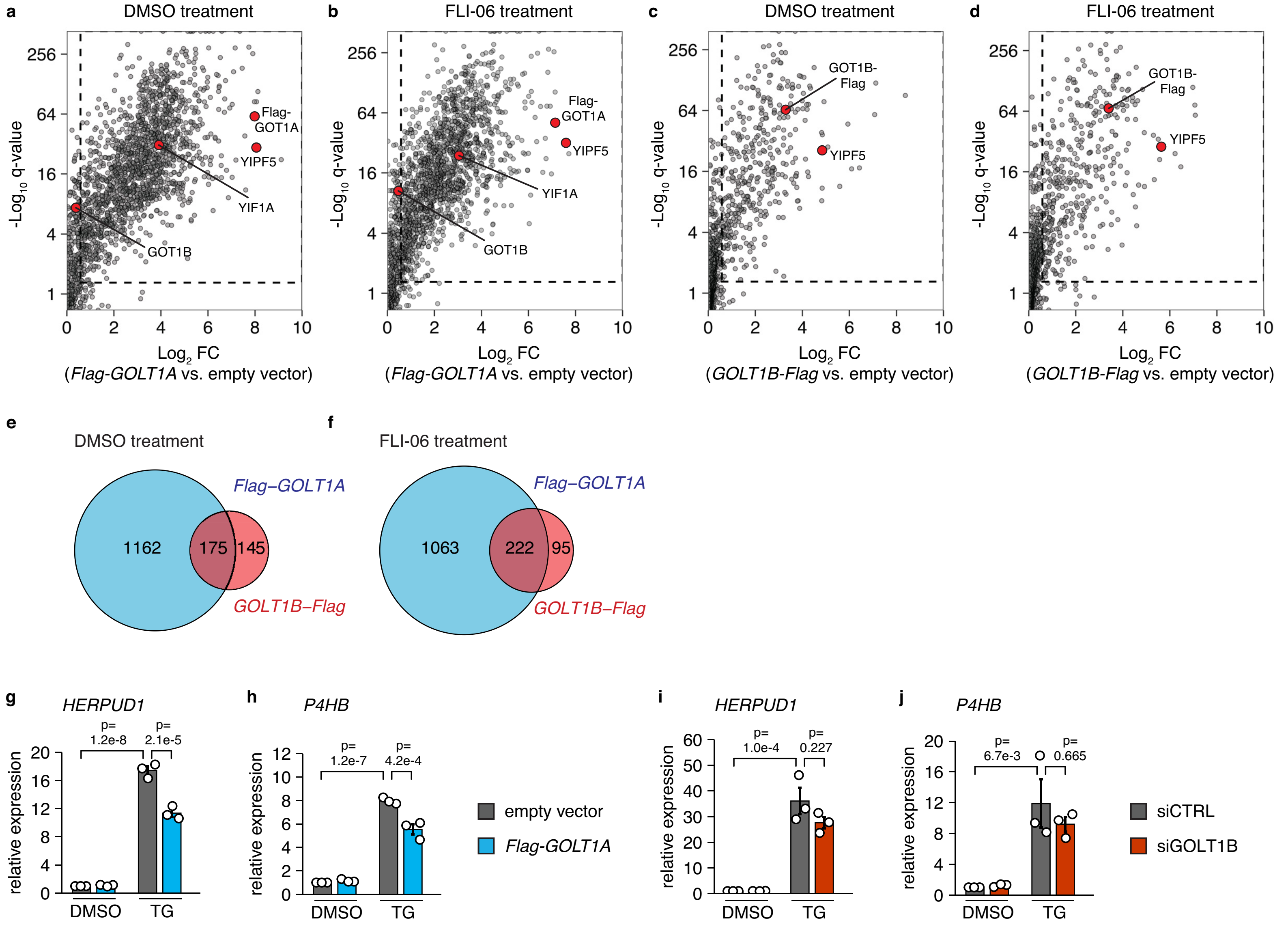
a
DMSO treatment
-Log10 q-value
Log2 FC
(Flag-GOLT1A vs. empty vector)
Flag-GOT1A
YIPF5
YIF1A
GOT1B
b
FLI-06 treatment
Flag-GOT1A
YIPF5
YIF1A
GOT1B
c
DMSO treatment
(GOLT1B-Flag vs. empty vector)
GOT1B-Flag
YIPF5
d
FLI-06 treatment
GOT1B-Flag
YIPF5
e
DMSO treatment
Flag–GOLT1A
1162
175
145
GOLT1B–Flag
f
FLI-06 treatment
Flag–GOLT1A
1063
222
95
GOLT1B–Flag
g
HERPUD1
p= 1.2e-8
p= 2.1e-5
relative expression
DMSO
TG
h
P4HB
p= 1.2e-7
p= 4.2e-4
empty vector
Flag-GOLT1A
i
HERPUD1
p= 1.0e-4
p= 0.227
j
P4HB
p= 6.7e-3
p= 0.665
siCTRL
siGOLT1B

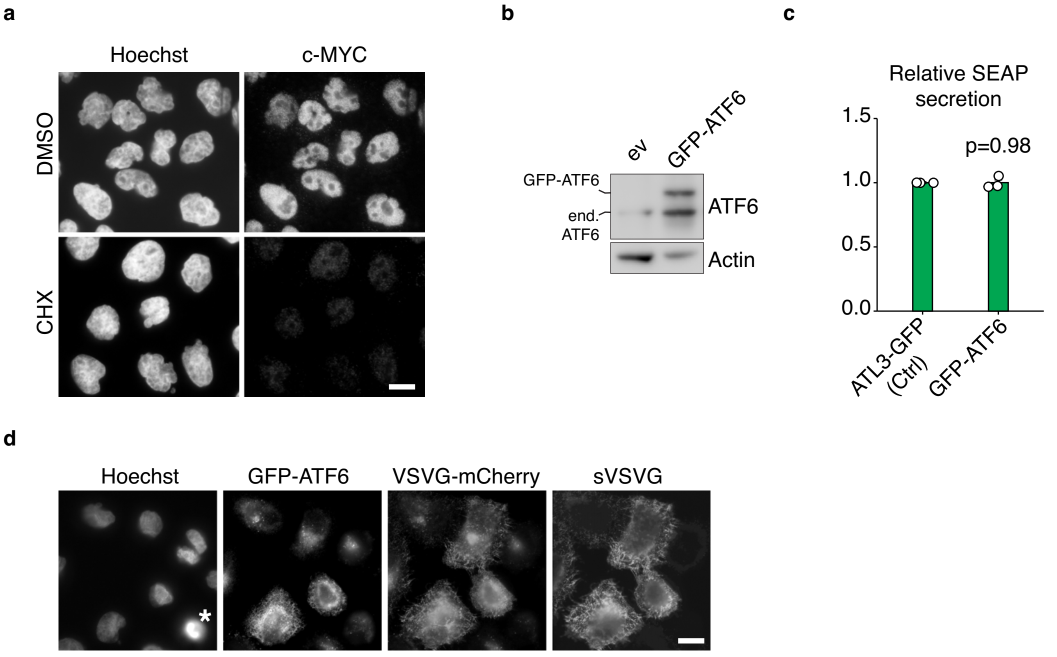
a
Hoechst
c-MYC
DMSO
CHX
b
ev
GFP-ATF6
GFP-ATF6
end.
ATF6
ATF6
Actin
c
Relative SEAP
secretion
1.5
1.0
0.5
0.0
p=0.98
ATL3-GFP
(Ctrl)
GFP-ATF6
d
Hoechst
GFP-ATF6
VSVG-mCherry
sVSVG
*

Supplemental experimental procedures

*Primary antibodies*

| Name | Application, dilution | Reference | RRID |
|---|---|---|---|
| anti-Actin (beta) | WB (1:5000) | Abcam, ab8227 | AB_2305186 |
| anti-bCOP | IF (1:500) | Invitrogen, PA1-061 | AB_2081296 |
| anti-ERGIC53 | IF (1:250) | Santa Cruz, sc-271517 | AB_10649805 |
| anti-FlagM2 | WB (1:1000), IF (1:500) | Sigma-Aldrich, F1804 | AB_262044 |
| anti-GFP | IF (1:2000) | Abcam, ab13970 | AB_300798 |
| anti-GolginB1 | IF (1:500) | Sigma-Aldrich, HPA011008 | AB_1079011 |
| anti-GOT1A | WB (1:1000) | Custom-made antibody provided by Eurogentec (clone #1870) | --- |
| anti-GOT1B | WB (1:1000) | Thermo Fisher Scientific, PA5-68055 | AB_2691673 |
| anti-HA | IF (1:1000) | Biolegend, 901502 | AB_2565007 |
| anti-RTN4 | IF (1:200) | Abcam, ab47085 | AB_881718 |
| anti-SEC16A | IF (1:200) | Abcam, ab70722 | AB_1270588 |
| anti-Vinculin | WB (1:50’000) | Sigma-Aldrich, V9131 | AB_477629 |
| anti-VSVG | WB (1:1000) | Kera Fast, EB0010 | AB_2811223 |
| anti-V5 | WB (1:1000) | Cell Signaling, 13202S | AB_2687461 |
| anti-YIF1A | WB (1:1000) | (Yoshida et al., 2008) | --- |
| anti-YIPF5 | WB (1:500), IF (1:250) | Invitrogen, PA5-67301 | AB_2664322 |
| anti-ATF6 | WB (1:1000), C&R (1:100) | Abcam #ab227830 | --- |
| anti-XBP1 | C&R (1:100) | Cell Signaling #40435 | AB_2891025 |
| anti-c-myc | IF (1:200) | Abcam Y69 #ab32072 | AB_731658 |
| anti-mCherry | | Abcam #ab167453 | AB_2571870 |

*Secondary antibodies*

| Name | Application, dilution | Reference | RRID |
|---|---|---|---|
| anti-chicken AF488 | IF (1:1000) | Thermo Fisher, A-11039 | AB_142924 |
| anti-mouse AF564 | IF (1:1000) | Thermo Fisher, A-11018 | AB_2534085 |
| anti-mouse IgG1 AF564 | IF (1:1000) | Thermo Fisher, A-21123 | AB_141592 |
| anti-mouse IgG2a AF647 | IF (1:1000) | Thermo Fisher, A-21241 | AB_141698 |
| anti-rabbit AF647 | IF (1:1000) | Thermo Fisher, A-21244 | AB_2535812 |
| anti-rabbit IgG-HRP | (WB, 1:10’000) | Santa Cruz Biotechnology, sc-2357 | AB_628497 |
| mouse-IgGκ BP-HRP | (WB, 1:10’000) | Santa Cruz Biotechnology, sc-516102 | AB_2687626 |

*Plasmids/libraries*

| Name | Reference/addgene # | kindly provided by |
|---|---|---|
| lenti_sgRNA(MS2)_zeo_backbone | Addgene #61427, (Konermann et al., 2015) | Feng Zhang |
| SGEP | Addgene #111170, (Fellmann et al., 2013) | Johannes Zuber |

| | | |
|---|---|---|
| LeGO-iG2 | Addgene #27341, (Weber et al., 2008) | Boris Fehse |
| psPAX2 | Addgene #12260 | Didier Trono |
| pMD2.G | Addgene #12259 | Didier Trono |
| human SAM sgRNA library | Addgene #1000000057 | Feng Zhang |
| human GeCKO lentiviral sgRNA library v2 | Addgene #1000000048 | Feng Zhang |
| dCas9-VP64 | Addgene #61425, (Konermann et al., 2015) | Feng Zhang |
| MCP-p65-HSF1 | Addgene #61426, (Konermann et al., 2015) | Feng Zhang |
| pcDNA3.1_VSVG-EYFP_ts045 | (Yonemura et al., 2016) | - |
| pEGFP-ATF6 | Addgene #32955, (Chen et al., 2002) | Ron Prywes |
| pEGFP-C1-GFP-ATL3 | (Behrendt et al., 2021) | - |

ON TARGETplus SMARTpool siRNAs (Dharmacon)

| Target gene | Catalog number |
|---|---|
| CTRL (control) | D-001810-10 |
| ATF6 | L-009917-00-0005 |
| GOLT1B | L-015393-00 |
| YIPF5 | L-018962-01 |

*Oligonucleotides*

qPCR primer

| Target gene | Sequence of forward primer (5'-3') | Sequence of reverse primer (5'-3') |
|---|---|---|
| h*ATF3* | CCTCTGCGCTGGAATCAGTC | TTCTTTCTCGTCGCCTCTTTTT |
| h*B2M* | GTGCTCGCGCTACTCTCTC | GTCAACTTCAATGTCGGAT |
| h*EPHX1* | TGAGGAGATCCACGACTTACAC | CATTCCGCCAGTAGGAGATGA |
| h*GBF1* | AGTGATTCGCTCTGAAGATACCA | GTGCCCACAAAACGAGCAT |
| h*GOLT1A* | GGTGTGGTTATCGTGCTCCTA | CGAAGGCGACAGGGAAAAAG |
| h*GOLT1B* | TCTGGGTGGTGTATTTGTAGTCC | CCAACAACGACAGGAAAGAAGCC |
| h*GPR89B* | CCGACTACTGCATAAACAACGA | GGCTGAGAATGGGAAAGGGAT |
| h*HERPUD1* | TGCTGGTTCTAATCGGGGACA | CCAGGGGAAGAAAGGTTCCG |
| h*PCNX2* | AGGAGGCCAAAGGTGAACAC | GACAACTCCAATGGCCCAGA |
| h*PRRC1* | CCAGAGGAGCAAGAAGACCC | CCAGGGTCCAGCGTTGTAAT |
| h*P4HB* | GGTGCTGCGGAAAAGCAAC | ACCTGATCTCGGAACCTTCTG |
| h*RNF13* | ACCTCCCTGCAAGATTTGGTT | CACTATGGGTTCACAGGCATTC |
| h*TMEM52* | CAGCACTGTGACCTCCTACA | TTTCTGGGAGAGTGCTGGTC |

Cloning of sgRNA and shRNA

| name | Forward primer (5'-3') | Reverse primer (5'-3') |
|---|---|---|

| | | |
|---|---|---|
| SAM sgRNA h*ATF3* | CACCGGGTGTGTGTCTCAGTG AGCG | AAACCGCTCACTGAGACACA CACCC |
| SAM sgRNA h*GBF1 #1* | CACCGCCTAAAGAAGCGCGAT TACA | AAACTGTAATCGCGCTTCTTT AGGC |
| SAM sgRNA h*GBF1 #2* | CACCGGGGAAAGAAAGGGTAT GTTA | AAACTAACATACCCTTTCTTT CCCC |
| SAM sgRNA h*GOLT1A #1* | CACCGCGCGAAACTTAGATTG TATG | AAACCATACAATCTAAGTTTC GCGC |
| SAM sgRNA h*GOLT1A #2* | CACCGGAGAGCCGCAGAACAC CGCG | AAACCGCGGTGTTCTGCGGC TCTCC |
| SAM sgRNA h*GOLT1A #3* | CACCGGGGTGCTCAAGTTTCT TGCT | AAACAGCAAGAAACTTGAGC ACCCC |
| SAM sgRNA h*GOLT1B #1* | CACCGAAGATTTTACTCCCGA GTAG | AAACCTACTCGGGAGTAAAA TCTTC |
| SAM sgRNA h*GOLT1B #2* | CACCGGAGCATCCACCCTTCC GGGT | AAACACCCGGAAGGGTGGAT GCTCC |
| SAM sgRNA h*GOLT1B #3* | CACCGGGAAAGATCTGCTCGA GGCC | AAACGGCCTCGAGCAGATCT TTCCC |
| SAM sgRNA h*GPR89B* | CACCGGTGCTAAAAGCAGCAA TGCA | AAACTGCATTGCTGCTTTTAG CACC |
| SAM sgRNA h*EPHX1* | CACCGCCTGGCAGAGGTGGA GCCTT | AAACAAGGCTCCACCTCTGC CAGGC |
| SAM sgRNA NT #1 | CACCGCTGAAAAAGGAAGGAG TTGA | AAACTCAACTCCTTCCTTTTT CAGC |
| SAM sgRNA NT #2 | CACCGAAGATGAAAGGAAAGG CGTT | AAACAACGCCTTTCCTTTCAT CTTC |
| SAM sgRNA h*PCNX2* | CACCGGCGAAGGCTAAGGAG GGACT | AAACAGTCCCTCCTTAGCCT TCGCC |
| SAM sgRNA h*PRRC1* | CACCGCTCGGTAGTTAGGAAG ATCT | AAACAGATCTTCCTAACTACC GAGC |
| SAM sgRNA h*RNF13* | CACCGTTAGAGATGCGAGCGG CCCG | AAACCGGGCCGCTCGCATCT CTAAC |
| SAM sgRNA h*TMEM52* | CACCGCTGCGGCCAGCGGGG CCCGG | AAACCCGGGCCCCGCTGGC CGCAGC |
| mir-E shRNA amplification | TGAACTCGAGAAGGTATATTG CTGTTGACAGTGAGCG | TCTCGAATTCTAGCCCCTTG AAGTCCGAGGCAGTAGGC |
| mir-E shRNA template h*GOLT1B* | TGCTGTTGACAGTGAGCGCTA GTATTATTGAAGATACTAATAG TGAAGCCACAGATGTATTAGTA TCTTCAATAATACTATTGCCTA CTGCCTCGGA | --- |

Supplemental references

Behrendt L, Hoischen C, Kaether C (2021) Disease-causing mutated ATLASTIN 3 is excluded from distal axons and reduces axonal autophagy. Neurobiol Dis 155: 105400

Chen X, Shen J, Prywes R (2002) The luminal domain of ATF6 senses endoplasmic reticulum (ER) stress and causes translocation of ATF6 from the ER to the Golgi. J Biol Chem 277: 13045-52

Fellmann C, Hoffmann T, Sridhar V, Hopfgartner B, Muhar M, Roth M, Lai DY, Barbosa IA, Kwon JS, Guan Y, Sinha N, Zuber J (2013) An optimized microRNA backbone for effective single-copy RNAi. Cell Rep 5: 1704-13

Konermann S, Brigham MD, Trevino AE, Joung J, Abudayyeh OO, Barcena C, Hsu PD, Habib N, Gootenberg JS, Nishimasu H, Nureki O, Zhang F (2015) Genome-scale transcriptional activation by an engineered CRISPR-Cas9 complex. Nature 517: 583-8

Weber K, Bartsch U, Stocking C, Fehse B (2008) A multicolor panel of novel lentiviral "gene ontology" (LeGO) vectors for functional gene analysis. Mol Ther 16: 698-706

Yonemura Y, Li X, Muller K, Kramer A, Atigbire P, Mentrup T, Feuerhake T, Kroll T, Shomron O, Nohl R, Arndt HD, Hoischen C, Hemmerich P, Hirschberg K, Kaether C (2016) Inhibition of cargo export at ER exit sites and the trans-Golgi network by the secretion inhibitor FLI-06. J Cell Sci 129: 3868-3877

Yoshida Y, Suzuki K, Yamamoto A, Sakai N, Bando M, Tanimoto K, Yamaguchi Y, Sakaguchi T, Akhter H, Fujii G, Yoshimura S, Ogata S, Sohda M, Misumi Y, Nakamura N (2008) YIPF5 and YIF1A recycle between the ER and the Golgi apparatus and are involved in the maintenance of the Golgi structure. Exp Cell Res 314: 3427-43
